# Stage-specific metabolic rewiring coordinates nucleotide supply and demand during spermatogenesis

**DOI:** 10.64898/2026.03.12.711306

**Authors:** Guy B Paz, Nina Mayorek, Yousef Mansour, Ilan Stein, Eleonora Medvedev, Sharona Elgavish, Shmuel Ruppo, Oriya Vardi, Boris Sarvin, Nikita Sarvin, Tomer Shlomi, Eli Pikarsky

## Abstract

Male meiotic prophase I features a major transition from leptotene/zygotene to pachytene/diplotene characterized by a massive transcriptomic and proteomic shift, yet how metabolism is coordinated across this transition remains unclear. Here, using stage-resolved purification of spermatogenic cells combined with respirometry, proteomics, metabolomics, and isotope tracing, we define metabolic programs across these stages. Early prophase cells rely on glutamine and fatty acid β-oxidation to sustain mitochondrial respiration while directing glucose into the pentose phosphate pathway and *de novo* pyrimidine synthesis. In contrast, late prophase cells utilize glucose and lactate as fuels, suppress fatty-acid oxidation, lose pentose phosphate pathway entry, and silence nucleotide biosynthesis despite a surge in nascent RNA synthesis and lower nucleotide pools. Inhibiting nucleoside uptake does not impair meiotic progression, indicating that late prophase transcription depends primarily on nucleotide pools synthesized earlier. These findings reveal a developmentally programmed separation between nucleotide production in early prophase and nucleotide use in later meiotic stages.

**TEASER:** Meiotic germ cells rewire central carbon metabolism to match stage-specific nucleotide production and RNA demand during spermatogenesis.

## INTRODUCTION

Spermatogenesis and spermiogenesis are sequential developmental processes that culminate in the production of mature spermatozoa. Diploid spermatogonial stem cells (SSCs), located at the periphery of the seminiferous tubules, sustain male reproductive capacity throughout life. Spermatogenesis is a complex, multistage process initiated by retinoic acid (*1*, *2*), during which SSCs undergo successive mitotic divisions before differentiating into preleptotene primary spermatocytes that enter the premeiotic S phase. The final round of DNA replication occurs in preleptotene (preL) cells (*3*), after which primary spermatocytes maintain a 4C DNA content throughout meiosis I as they progress through meiotic prophase I (leptotene, zygotene, pachytene, and diplotene). Although recombination may involve short tracts of repair-associated DNA synthesis, no further bulk DNA replication occurs.

Importantly, spermatocytes progress sequentially from leptotene to zygotene, through early, mid, and late pachytene, and then to diplotene and diakinesis before completing the first meiotic division. Two consecutive meiotic divisions ultimately produce haploid round spermatids (RS), which undergo spermiogenesis to form mature spermatozoa. (*4*).

This developmental program takes place within the seminiferous tubules, where premeiotic cells at the periphery have direct access to nutrients from nearby blood vessels. Upon entry into meiosis at the preL stage, cells cross the blood-testis barrier (BTB) via Sertoli cell tight junctions to the adluminal compartment, where differentiation proceeds and nutrient supply is provided by Sertoli cells and the seminiferous tubule fluid (*5*).

In mice, spermatogenesis and spermiogenesis span ∼35 days and require substantial metabolic adaptation. Advances in methods for isolating cells at discrete stages have revealed a major transcriptomic switch between the leptotene/zygotene (LZ) and pachytene/diplotene (PD) stages (*6–8*). However, despite extensive transcriptional remodeling during this transition-affecting >10,000 genes(*9*) it remains unclear how these large-scale changes are translated into functional metabolic programs, that can support the dramatic increases in biomass and biosynthetic output. This gap is particularly striking given the rapid increase in cellular biomass across the same interval: cell volume increases ∼3.4-fold between LZ and PD in prepubertal mice(*10*) and ∼5-fold in adults (*9*), accompanied by a ∼4-fold increase in RNA and protein content, implying substantial bioenergetic and biosynthetic remodeling.

Here, we systematically examine the metabolic challenges faced by developing germ cells and identify a metabolic switch between the LZ and PD stages, characterized by coordinated changes in the metabolome, mitochondrial content and activity, and fuel preferences. Using ^13^C-glucose tracing, we find that LZ cells channel glucose away from energy production and toward anabolic pathways that support nucleotide synthesis, including the pentose phosphate pathway (PPP). In contrast, PD cells lose PPP activity coincident with transcriptional silencing of glucose-6-phosphate dehydrogenase (G6PD), an X-linked enzyme, as a consequence of meiotic sex chromosome inactivation (MSCI) (*11*). In parallel, we observe downregulation of rate-limiting enzymes involved in *de novo* purine and pyrimidine synthesis as cells transition from LZ to PD.

Together, these findings support a model in which meiotic metabolism is organized around a developmentally enforced separation between nucleotide production and nucleotide utilization: LZ cells generate nucleotide pools that support subsequent stages, whereas PD cells, despite continued biosynthetic activity, show negligible capacity for *de novo* nucleotide synthesis. We further test the contribution of extracellular nucleoside uptake using the ENT1 inhibitor, NBMPR (*12*) and find that differentiating germ cells rely predominantly on internal nucleotide sources. Finally, comparative analysis across mouse, human, zebrafish, and yeast indicates that repression of nucleotide biosynthetic gene expression during late meiotic prophase I is evolutionarily conserved, spanning vertebrate spermatogenesis and sporulation-associated meiosis in *S. cerevisiae*.

## RESULTS

### Metabolome profiling of spermatogenic cells reveals a metabolic switch between LZ and PD populations

As described above, a profound shift in gene expression occurs during prophase I of spermatogenesis, particularly between the LZ and PD populations. To examine how these transcriptional changes shape cellular metabolism, we isolated discrete germ-cell populations spanning spermatogonia (Spg) to RS using our previously developed flow-cytometry method **(Fig. S1)** (*9*) and subjected them to LC-MS analysis.

LC-MS profiling identified 108 metabolites across 4 to 5 biological replicates per population. Given the substantial changes in germ cell volume across prophase I (*9*), metabolite abundances were normalized to cell volume to account for stage-dependent differences in cell size and to better approximate relative intracellular metabolite concentrations. 70 metabolites were differentially expressed among all populations (Kruskal-Wallis test, adjusted p < 0.05) (**Table S1**). Principal component analysis (PCA) revealed tight clustering of replicates within each cell type, indicating a distinct metabolic profile for each developmental stage (**Fig. 1A**). Notably, PC1 captured ∼44% of total variance and arranged cell populations along their biological developmental order, separating Spg-LZ from PD-RS and revealing a coordinated, stage-aligned metabolic reprogramming during meiosis.

**Fig. 1.**
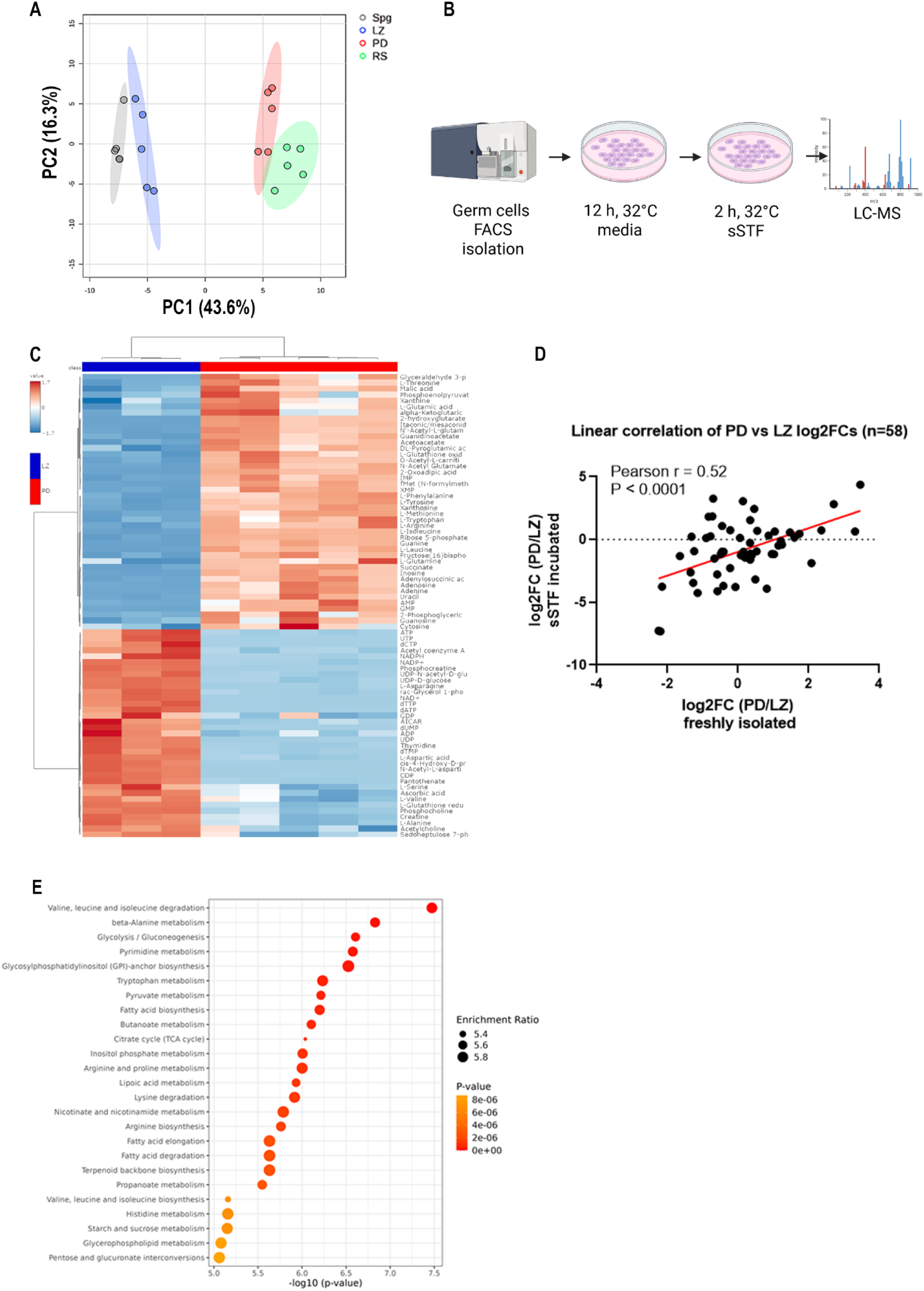
Metabolome profiling of spermatogenic cells reveals a metabolic switch between LZ and PD populations. (**A**) PCA of LC-MS metabolomics data from freshly isolated germ-cell populations: spermatogonia (Spg), leptotene/zygotene (LZ), pachytene/diplotene (PD), and round spermatids (RS) (n = 4-5 biological replicates per population). (**B**) Workflow for metabolomics of *ex vivo*- recovered LZ and PD cells: cells were isolated by FACS, incubated for 12 h at 32°C in MEM medium, followed by 2h incubation in synthetic seminiferous tubule fluid (sSTF) prior to metabolite extraction and LC-MS analysis. This figure was created with BioRender.com. (**C**) Heat map showing metabolites with significant differences across groups (Kruskal-Wallis test, adjusted p < 0.05) in overnight-recovered and sSTF-incubated LZ (n = 3) and PD (n = 5) cells. For each sample, the color intensity-from blue to red-represents the z-score of the corresponding metabolite, as indicated by the scale in the figure. **(D)** Correlation of PD/LZ ratio of metabolites extracted from freshly isolated (Table S1) and *ex vivo*-incubated (Table S3) spermatogenic cells. Scatter plot shows log₂ fold change (PD/LZ) for each shared metabolite measured in freshly isolated cells (x-axis) versus sSTF-incubated cells (y-axis). A total of 56 co-detected metabolites were included. Linear regression revealed a moderate positive correlation between datasets (Pearson r=∼0.52, P < 0.0001), indicating partial conservation of the direction and magnitude of metabolic changes between LZ and PD across experimental conditions. (**E**) KEGG pathway enrichment analysis of metabolites differentially abundant between LZ and PD cells incubated in sSTF (dot size indicates enrichment ratio; color indicates P value).

Fluorescence-activated cell sorting (FACS) can cause sorter-induced cellular stress, characterized by oxidative/redox perturbations, altered energy charge, and broad changes in metabolite abundances (*13*, *14*). In addition, cellular metabolomes are influenced by the composition of the incubation medium (*15–17*). Therefore, we examined the metabolomes of LZ and PD populations under conditions designed to approximate the *in vivo* environment. Sorted cells were incubated for 12 h in MEM medium (*9*) and then transferred to synthetic seminiferous tubule fluid (sSTF; **Table S2; Fig. 1B**). The sSTF formulation was based on previously published measurements of seminiferous tubule fluid composition (*18–21*).

This dataset comprised 81 detected metabolites, 76 of which differed significantly between LZ and PD (Kruskal-Wallis test, adjusted p < 0.05; **Table S3; Fig. 1C**), reaffirming extensive metabolic divergence between these stages. Comparing the two datasets (**Tables S1 and S3**) revealed that, of the 131 metabolites detected overall, 56 were shared between freshly isolated cells and sSTF-incubated cells (r = ∼0.52, p < 0.0001, Pearson correlation, comparing the changes between PD-versus-LZ cells between the two conditions; **Fig. 1D**). Together, these results demonstrate pronounced metabolic reprogramming as cells progress through meiotic prophase.

To identify the biological pathways driving this metabolic shift, we conducted KEGG-based pathway enrichment analysis of metabolites differentially expressed between LZ and PD incubated in sSTF medium. Most changes mapped to amino-acid metabolism, glycolysis, and pyrimidine metabolism, implicating central-carbon and nucleotide metabolic pathways as key features of the LZ-to-PD transition (**Fig. 1E**).

### LZ-to-PD transition is associated with significant mitochondrial reorganization

Mitochondria are central hubs for energy metabolism and biosynthetic integration. To better understand metabolic reprogramming across the LZ-to-PD transition(*9*), we examined mitochondrial protein composition using our previously generated proteomic dataset. Our analysis identified 3,916 proteins, of which 2,229 were significantly differentially expressed (**Fig. 2A**). By intersecting these significantly changing proteins with the MitoCarta 3.0 inventory (*22*), which comprises 1,140 experimentally validated mouse mitochondrial proteins, we identified 569 mitochondrial proteins expressed in LZ and/or PD cells, of which 243 were significantly differentially expressed between LZ and PD spermatocytes (**Fig. 2B**).

**Fig. 2.**
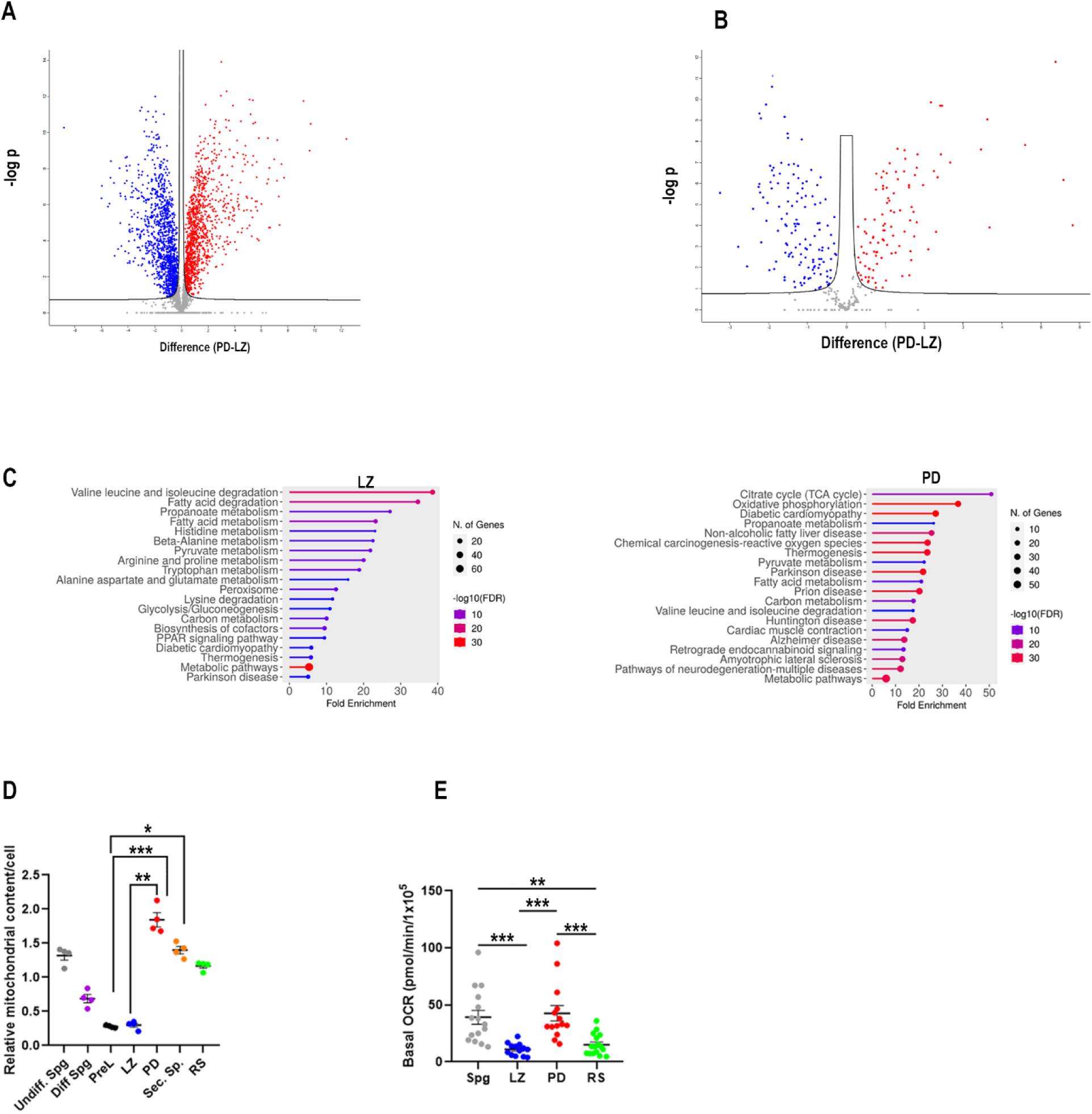
LZ-to-PD transition is associated with significant mitochondrial reorganization. (**A**) Volcano plot of proteomics data comparing protein abundance between PD and LZ spermatocytes (x-axis: abundance difference, PD-LZ; y-axis: −log P). Proteins significantly increased in PD or LZ are highlighted in red and blue, respectively (T-test, FDR-adjusted p < 0.05). (**B**) Volcano plot as in (A), restricted to mitochondria-associated proteins detected in our dataset based on intersection with the MitoCarta 3.0 inventory, highlighting mitochondrial proteins differentially abundant between LZ and PD. (**C**) KEGG pathway enrichment analysis of mitochondrial proteins enriched in LZ (left) or PD (right). The x-axis indicates fold enrichment; dot size reflects the number of proteins in each pathway; color indicates statistical significance (-log_10_ FDR). analysis was performed using Enrichr web tool.(*23*) (**D**) Relative mitochondrial content per cell across spermatogenic populations measured by MitoTracker Green staining (undifferentiated spermatogonia, differentiating spermatogonia, preleptotene, LZ, PD, and round spermatids). Points represent experimental replicates (n=4); horizontal bars indicate mean ± SEM; P values are shown (ANOVA test). (**E**) Basal oxygen consumption rate (OCR) measured by Seahorse respirometry in the indicated germ-cell populations. Points represent individual samples (n=13-16); horizontal bars indicate mean ± SEM; (ANOVA test). *P < 0.05; **P < 0.01; ***P < 0.001.

Pathway analysis of mitochondrial proteins enriched in LZ cells indicated active amino acid metabolism and fatty acid β-oxidation. In contrast, mitochondrial proteins enriched in PD cells were predominantly associated with the tricarboxylic acid (TCA) cycle and oxidative phosphorylation (OX-PHOS) (**Fig. 2C**). These data suggest a functional mitochondrial reorganization during the LZ-to-PD transition, characterized by a shift from amino acid and fatty acid catabolism toward enhanced energy production via the TCA cycle and oxidative phosphorylation.

Next, we assessed mitochondrial content across germ-cell populations during spermatogenesis using MitoTracker Green staining. Mitochondrial content decreased progressively from Spg to PreL and LZ stages, followed by a significant elevation in PD cells **(Fig. 2D, Fig S2**). This trend was corroborated by Seahorse analysis, which revealed a ∼3.5-fold decrease in basal oxygen consumption rate (OCR) from Spg to LZ, followed by a ∼4-fold increase from LZ to PD (**Fig. 2E).** Collectively, these results indicate that spermatocytes undergo both quantitative and functional mitochondrial reorganization during the LZ-to-PD transition.

### LZ-to-PD transition is associated with a shift in fuel utilization preferences

Motivated by the observed changes in mitochondrial functional capacity, we investigated substrate preference in LZ and PD spermatocytes using Seahorse respirometry. The addition of 5 mM glucose or lactate to LZ cells incubated in basal buffer lacking carbon sources did not alter oxygen consumption rate (OCR). In contrast, PD cells exhibited ∼1.7-and ∼3.0-fold increases in OCR following addition of 5 mM glucose or lactate, respectively, under the same conditions (**Fig. 3A**; **Fig. S3A**). Notably, Spg exhibited a negligible response to lactate, similar to LZ cells, whereas RS showed a response comparable to that of PD cells. (**Fig S3B**).

**Fig. 3.**
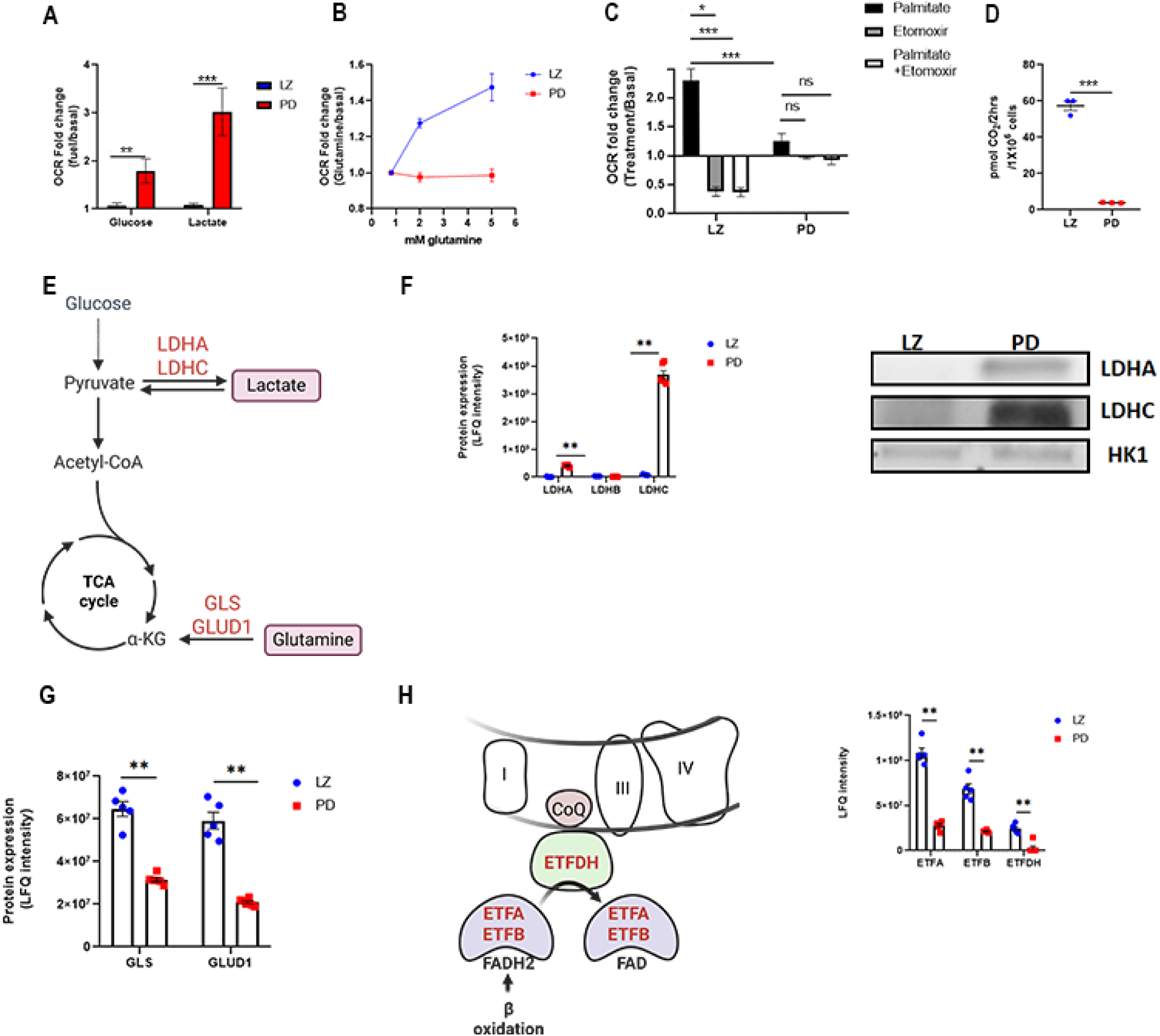
LZ-to-PD transition is associated with a shift in fuel utilization preferences. (**A**) Seahorse respirometry showing the change in OCR (fold change relative to basal) in LZ and PD spermatocytes following addition of 5 mM glucose or lactate to cells incubated in carbon-free basal buffer. (Student T-test) **(B**) OCR response (fold change relative to basal) of LZ and PD spermatocytes to increasing concentrations of glutamine. (**C**) OCR response (fold change relative to basal) following addition of 0.3 mM palmitate, the CPT1 inhibitor etomoxir 40 µM, or their combination, revealing CPT1-dependent fatty-acid oxidation in LZ but not PD cells (ANOVA). (**D**) Fatty-acid oxidation measured by ^14^CO_2_ production from ^14^C-palmitate in LZ and PD cells, see details in Methods section (Student T-test). (**E**) Scheme summarizing entry points of lactate and glutamine into glycolysis and TCA, respectively. This figure was created with BioRender.com. (**F**) Left: proteomics-based quantification (LFQ intensity) of LDHA, LDHB and LDHC in LZ and PD cells (Student T-test). Right: representative immunoblot validating LDHA and LDHC differences (HK1 shown as a control). (**G**) Proteomics-based quantification (LFQ intensity) of glutamine catabolic enzymes (GLS and GLUD1) in LZ and PD cells (Student T-test). (**H**) Left: scheme of electron transfer from β-oxidation to the respiratory chain via ETFA/ETFB and ETFDH to coenzyme Q (CoQ), enabling FAD regeneration and sustained β-oxidation. Right: proteomics-based quantification (LFQ intensity) of ETFA, ETFB, and ETFDH (Student T-test). For all plots, points represent biological replicates; bars/lines indicate mean ± SEM; *P < 0.05; **P < 0.01; ***P < 0.001; n.s., not significant

To identify the energy sources supporting LZ cells given their lack of response to glucose and lactate, we next tested glutamine as a potential substrate. LZ, but not PD cells, showed increased OCR upon glutamine supplementation, with ∼1.3-and ∼1.6-fold increases following addition of 2 mM or 5 mM glutamine, respectively (**Fig. 3B**).

Strikingly, consistent with the enrichment of mitochondria-localized protein pathways (**Fig. 2C**), fatty acids emerged as a major energy source in LZ cells. Palmitate increased OCR by ∼2.3-fold in LZ cells but had little to no effect on basal OCR in PD cells. Moreover, etomoxir-an irreversible inhibitor of carnitine palmitoyltransferase 1 (CPT1)-reduced OCR in LZ cells by ∼60%, with no significant change in PD cells. Co-treatment with palmitate and etomoxir recapitulated the effect of etomoxir alone, supporting the specificity of CPT1-dependent fatty acid oxidation in LZ cells **(Fig. 3C**). These findings were further validated by measuring ^14^CO_2_ production from ^14^C-palmitate, which demonstrated active fatty acid oxidation in LZ cells, but no detectable palmitate utilization via this pathway in PD cells (**Fig. 3D**).

Our proteomic data provide mechanistic insight into these metabolic differences. The inability of LZ cells to utilize lactate is consistent with markedly reduced expression of LDHA and LDHC. LDHB was expressed at negligible levels in both cell populations. **(Fig. 3E, F)**. The lack of glutamine utilization by PD cells aligns with reduced expression of glutamine catabolic enzymes, including GLS and GLUD1 (**Fig. 3E, G**).

Finally, the lack of fatty acid oxidation in PD cells is attributable to the absence of ETFDH and markedly reduced levels of ETFA and ETFB, which are required to reoxidize FADH_2_ and sustain β-oxidation. FADH_2_ is generated by mitochondrial acyl-CoA dehydrogenases (ACADs), and electrons are transferred via ETF/ETFDH to coenzyme Q in the electron transport chain, regenerating FAD and enabling continued β-oxidation (*24*) (**Fig. 3H**).

Together, these data reveal a shift in fuel utilization during spermatocyte differentiation, whereby LZ cells preferentially rely on fatty acid oxidation and glutamine metabolism, while PD and RS cells shift toward glucose and lactate utilization.

### Differential metabolic fates of glucose in LZ and PD cells

The unexpected absence of an OCR response to glucose addition in LZ cells prompted us to investigate the metabolic fate of glucose in spermatocytes. We subjected both LZ and PD populations to [U-^13^C]-glucose tracing analysis. Cells were allowed to recover from FACS over-night prior to 2 h tracing in sSTF media. Experimental design, along with schematic representation of glucose tracing into glycolysis and pentose phosphate pathway (PPP) metabolites is depicted in (**Fig 4A**).

**Fig. 4.**
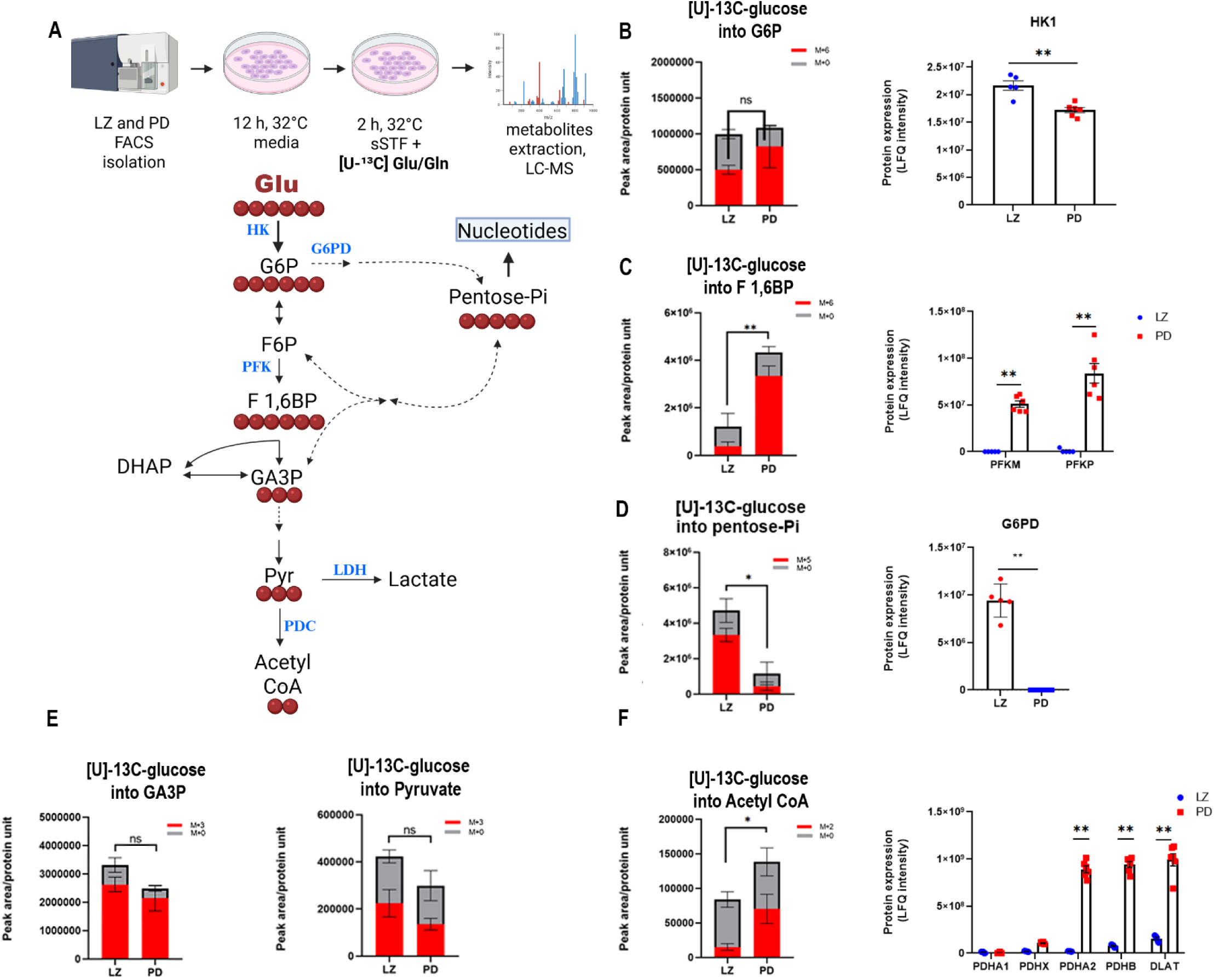
Differential metabolic fates of glucose in LZ and PD cells. (**A**) Sorted cells were recovered over-night and then incubated for 2 h in sSTF containing [U-¹³C]-glucose. The scheme illustrates labeling of glycolysis and PPP intermediates from [U-¹³C]-glucose; each red ball represents a carbon atom derived from the labeled glucose. This figure was created with BioRender.com. (**B**) Left: total and M+6-labeled glucose-6-phosphate (G6P) in LZ and PD cells, right: protein abundance (LFQ) of hexokinase 1 (HK1). (**C**) Left: total and M+6-fructose 1,6 bisphosphate (F1,6BP) in LZ and PD cells, right: protein abundance (LFQ) of phosphofructokinase isoforms M and P (PFKM and PFKP) (**D**) PPP activity is present in LZ but absent in PD cells. Left: total and M+5 labeling of pentose-5-phosphate. Note that the LC-MS method used does not resolve individual pentose phosphate isomers (e.g., ribose-5-P/ribulose-5-P/xylulose-5-P). Right: protein abundance (LFQ) of G6PD, the rate-limiting enzyme for PPP entry. (E) Re-entry of PPP-derived carbon into lower glycolysis: total and M+3 labeling of glyceraldehyde-3-phosphate (GA3P) and pyruvate in LZ and PD cells.(F) Divergence at the pyruvate dehydrogenase complex (PDC): total and M+2 labeling of acetyl-CoA from glucose-derived pyruvate is low in LZ compared with PD cells. Protein abundance (LFQ) of PDC components shows absent expression of multiple PDC subunits in LZ, whereas PDC subunits are highly expressed in PD. In PD cells, the X-linked PDHA1 isoform is absent (consistent with MSCI), whereas the autosomal testis-specific PDHA2 isoform is induced. Bars show mean ± SEM (n=3-4 independent [U-¹³C]-glucose tracing experiments). Proteomic enzyme abundance (mean ± SEM) was calculated from the PXD039015 dataset; points denote 5 biological replicates for LZ and 6 biological replicates for PD. For all labeling and proteomic quantifications, statistical significance was assessed using a Student’s T-test (*P < 0.05; **P < 0.01; n.s., not significant).

This analysis revealed that the levels of glucose 6-phosphate (G6P), both M+6-labeled and unlabled, were only slightly and insignificantly elevated in PD cells compared to LZ cells. In addition, hexokinase (HK1) protein expression-the enzyme responsible for glucose phosphorylation-was comparable between the two populations, suggesting that glucose uptake was not impaired in either cell type (**Fig. 4B**). However, the metabolic fate of G6P in LZ and PD cells was notably different. LZ cells were unable to proceed through the glycolytic pathway, as evidenced by their negligible M+6 tracing in fructose 1,6-bisphosphate (F1,6BP), which was ∼6-fold lower than that observed in PD cells. This surprising result was explained by the lack of expression of both phosphofructokinase isoforms (PFKM and PFKP) in LZ as compared to PD (**Fig. 4C**). Instead of entering glycolysis, G6P was redirected into the PPP. This was evidenced by the pronounced M+5 tracing of pentose-5 phosphates in LZ cells, which was not observed in PD cells. Furthermore, glucose 6-phosphate dehydrogenase (G6PD), the first enzyme that catalyzes entry into the PPP, was highly expressed at the protein level in LZ but was completely absent in PD cells (**Fig. 4D**). This absence is not unexpected, as G6PD is X-linked, and the transcription of chromosome X is silenced in PD cells due MSCI (*11*).

The glucose carbons that bypass the PFK step by entering PPP are reintegrated into glycolysis at the three-carbon level. No significant differences were observed in the M+3 tracing of glyceraldehyde 3-phosphate (GA3P) and pyruvate between the two cell populations (**Fig. 4E**).

The next metabolic divergence between LZ and PD cells was identified at the pyruvate dehydrogenase complex (PDC) step, that leads to acetyl-CoA formation. While PD cells proceed with the decarboxylation of glucose-derived pyruvate, LZ cells do not, as evidenced by M+2 tracing of acetyl CoA, which was ∼5-fold lower in LZ cells than in PD cells. Proteomic analysis revealed that expression of key PDC components was significantly higher in PD cells compared to LZ cells. Specifically, PDHA2, PDHB, and DLAT showed marked upregulation. (**Fig. 4F**). Notably, PDHA1, the X-linked isoform of the E1α subunit, was absent in PD cells, consistent with MSCI. To compensate, transcription of the autosomal, testis-specific PDHA2 isoform from chromosome 3 is activated, allowing continued PDC function in late prophase 1 cells (*25*). This limited flux of glucose-derived carbons into the TCA cycle in LZ cells may account for their inability to elevate OCR in response to glucose addition.

Together, these data demonstrate that glucose is differentially utilized in LZ and PD cells: while PD cells efficiently channel glucose through glycolysis and into the TCA cycle, LZ cells are metabolically restricted at key enzymatic steps, diverting glucose into the pentose phosphate pathway and limiting its contribution to oxidative metabolism.

### LZ cells, but not PD cells, are actively engaged in *de novo* pyrimidine nucleotide synthesis

Enhanced diversion of glucose into the PPP (**Fig 4D**), suggested that LZ cells engage in nucleotide biosynthesis, while PD cells may not do so. To test this directly, we traced [U-¹³C]-glucose and [U-¹³C]-glutamine incorporation into pyrimidine nucleotides as described in the experimental setup above (**Fig.5A)**.

**Fig. 5.**
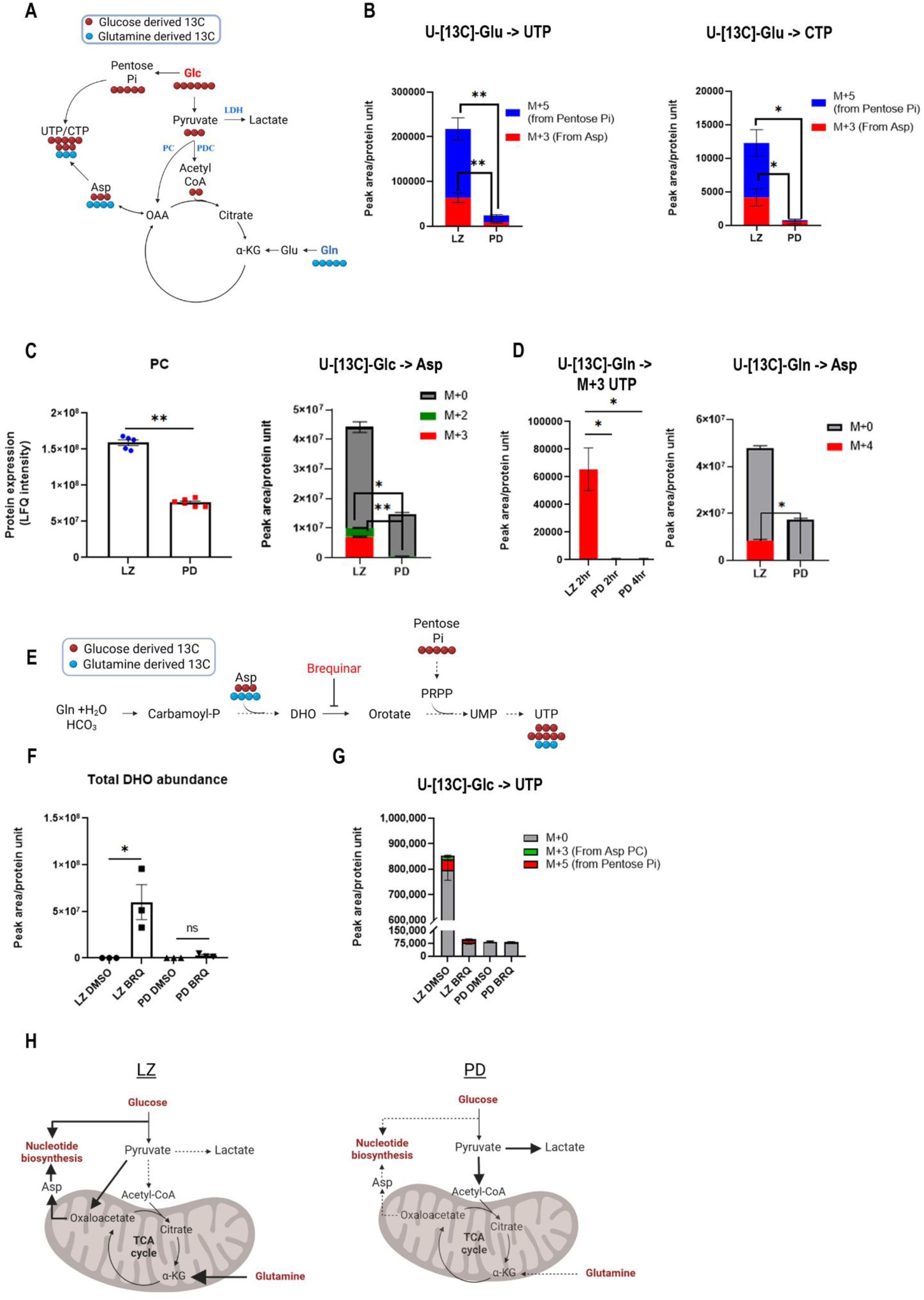
LZ cells, but not PD cells, are actively engaged in *de novo* pyrimidine nucleotide synthesis. **(A)** Scheme of isotope incorporation from [U-¹³C]-glucose and [U-¹³C]-glutamine into pyrimidine nucleotides. Glucose-derived carbon contributes to the ribose via the PPP (yielding M+5 nucleotides) and to the pyrimidine ring via pyruvate carboxylase (PC)**-**dependent anaplerosis (pyruvate → oxaloacetate → aspartate; yielding M+3 nucleotides). **(B)** Mass isotopologue labeling of UTP and CTP after [U-¹³C]-glucose tracing in LZ and PD cells. Stacked bars indicate M+5 (PPP-derived ribose; blue) and M+3 (aspartate-derived ring carbon; red). LZ cells show robust M+5 and M+3 labeling, whereas PD cells show little to no labeling. **(C)** Left: PC protein abundance in LZ and PD cells. Right: aspartate isotopologues after [U-¹³C]-glucose tracing (M+0, gray; M+2, green; M+3, red), showing prominent M+3 (and minor M+2) aspartate labeling in LZ cells but not PD cells. **(D)** Contribution of glutamine to pyrimidine-ring synthesis. Left: M+3 UTP after [U-¹³C]-glutamine tracing in LZ cells (2 h) and PD cells (2 and 4 h). Right: aspartate isotopologues after [U-¹³C]-glutamine tracing (M+0, gray; M+4, red), consistent with glutamine entry into the TCA cycle to generate labeled aspartate for incorporation into pyrimidines in LZ cells. **(E)** Scheme highlighting *de novo* pyrimidine synthesis pathway and inhibition of dihydroorotate dehydrogenase (DHODH) by brequinar (BRQ) at the dihydroorotate (DHO) → orotate step**. (F)** Total DHO abundance following DMSO or BRQ treatment in LZ and PD cells, showing BRQ-dependent DHO accumulation in LZ cells but not PD cells. **(G)** UTP isotopologues after [U-¹³C]-glucose tracing in DMSO- or BRQ-treated cells, showing reduction of unlabeled (M+0) and labeled (M+3 and M+5) UTP in BRQ-treated LZ cells. **(H)** Working model: LZ cells support *de novo* pyrimidine synthesis through PPP-derived ribose and PC/TCA cycle-supported aspartate production, with glutamine contributing via TCA cycle-derived aspartate; PD cells show limited routing of glucose/glutamine carbon into pyrimidine nucleotides. All tracing experiments were performed as shown in Fig 4A, cells were incubated for 2h unless stated otherwise. Bars show mean ± SEM (n =3-4 [U-¹³C]-glucose/glutamine tracing experiments). For all labeling and proteomic quantifications, statistical significance was assessed using a Student’s T-test (*P < 0.05; **P < 0.01). All schemes were created with BioRender.com.

In LZ cells, [U-¹³C]-glucose produced robust M+5 and M+3 labeling in UTP and CTP, whereas PD cells showed little to no labeling (**Fig. 5B**). The M+5 isotopologues reflect incorporation of glucose-derived carbons into the ribose moiety via the PPP, consistent with pentose phosphate labeling. The M+3 isotopologues are explained by glucose-derived pyruvate entering TCA cycle through pyruvate carboxylase (PC). PC protein expression was ∼2-fold higher in LZ than PD, generating labeled oxaloacetate (OAA) and subsequently aspartate. Consistent with this route, aspartate in LZ cells showed prominent M+3 labeling (and minor M+2), whereas PD cells did not (**Fig. 5C**). Because aspartate contributes three carbons to the pyrimidine ring, this accounts for the M+3 labeling observed in UTP and CTP (**Fig. 5A**).

The minor M+2 aspartate labeling from [U-¹³C]-glucose in LZ cells is consistent with partial loss of labeled carbon during TCA-cycle decarboxylation steps after OAA formation. This interpretation aligns with an active TCA cycle in LZ cells, as indicated by increased OCR after glutamine addition (**Fig. 3B**). Notably, because pyruvate does not generate acetyl-CoA in LZ cells due to an inactive PDC, acetyl-CoA is likely supplied by alternative fuels such as fatty acids via β-oxidation (**Fig. 3C, D**).

We also observed that glutamine contributes to pyrimidine-ring carbon in LZ cells but not PD cells. After [U-¹³C]-glutamine tracing, LZ cells exhibited M+4 aspartate accompanied by M+3 UTP labeling, consistent with glutamine feeding the TCA cycle to generate labeled aspartate that is then incorporated into pyrimidines (**Fig. 5D**).

We functionally corroborated the isotope-tracing results by pharmacologically inhibiting dihydroorotate dehydrogenase (DHODH) using brequinar (BRQ), a key rate-limiting enzyme in pyrimidine biosynthesis. (**Fig. 5E**) (*26*). Remarkably, BRQ treatment led to a marked accumulation of dihydroorotate (DHO) and depletion of UTP in LZ cells, whereas PD cells showed neither DHO accumulation nor UTP depletion. (**Fig. 5F, G**). Together, the absence of glucose-or glutamine-derived labeling in UTP and CTP in PD cells and the lack of BRQ-induced DHO accumulation strongly support that *de novo* pyrimidine synthesis is largely confined to the LZ stage (**Fig. 5H**).

### LZ-to-PD transition is associated with a surge in transcriptional activity

Nucleotide biosynthesis is a metabolically demanding process that requires substantial cellular resources (*27*). Our observations reveal a striking phenomenon: LZ cells channel central carbon metabolism toward robust *de novo* nucleotide synthesis, whereas these pathways are largely silenced in PD cells. This shift is particularly notable given the significant ∼5-fold increase in cell volume reported during the LZ-to-PD transition (*9*) and the parallel rise of ∼4-fold in protein content (**Fig 6A**). Such an expansion in cellular biomass requires a substantial increase in translational capacity.

**Fig. 6.**
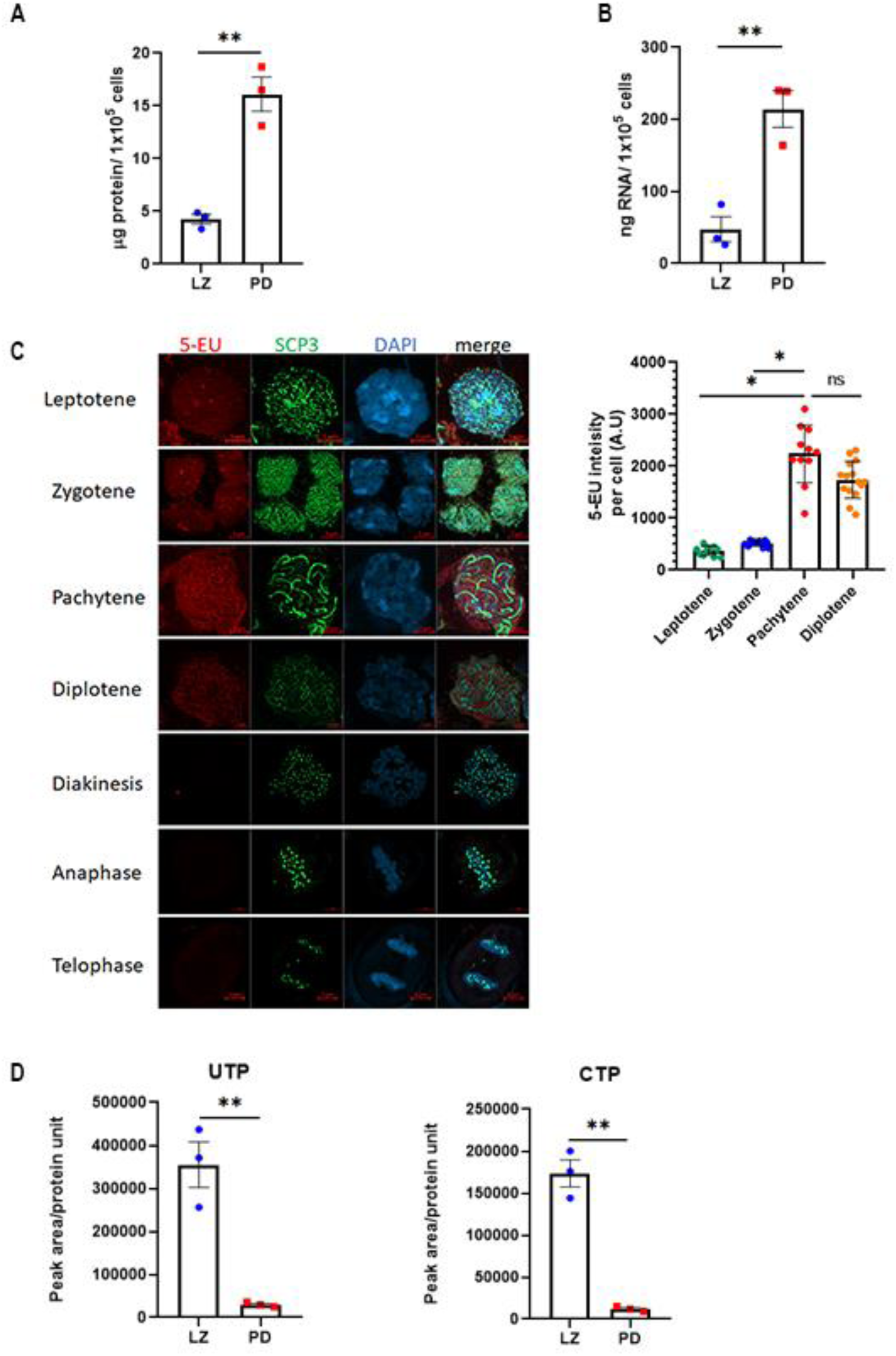
LZ-to-PD transition is associated with a surge in transcriptional activity and increased nucleotide utilization. **(A)** Total protein content per cell in LZ and PD populations, showing increased protein abundance in PD cells (normalized per 1 × 10⁵ cells), n=3 independent experiments (Student’s T-test). **(B)** Total RNA content per cell in LZ and PD populations, showing increased RNA abundance in PD cells (normalized per 1 × 10⁵ cells), n=3 independent experiments (Student’s T-test). **(C)** Left: nascent RNA synthesis measured by 5’-ethynyl uridine (5-EU) incorporation during a 2-h period. Representative images of meiotic stages (leptotene, zygotene, pachytene, diplotene, diakinesis, anaphase, telophase) showing 5-EU signal (red), SCP3 (green), and DNA (DAPI, blue). Right: quantification of 5-EU fluorescence intensity per cell across prophase I stages. Bars show mean ± SEM (n =12-15 cells), (ANOVA) **(D)** Relative abundance of pyrimidine nucleotides UTP and CTP in LZ and PD cells (peak area normalized to protein), showing reduced nucleotide pools in PD cells compared with LZ cells. Bars show mean ± SEM (n =3 independent experiments) (Student’s T-test). *P < 0.05; **P < 0.01 n.s., not significant for all graphs.

Consistent with this expectation, we observed a ∼4.5-fold increase in total RNA content accompanying the LZ-to-PD transition (**Fig 6B**), indicating enhanced ribosome biogenesis. Previous studies similarly reported elevated mRNA synthesis rates in PD cells relative to LZ cells (*28-29*).

To quantify transcriptional activity in our system, we measured the incorporation of 5’-ethynyl-uridine (5-EU) into nascent RNA over a 2 h labeling period across distinct meiotic populations (**Fig 6C**). Using click chemistry to generate fluorescent readouts and microscopic imaging for quantification, we characterized transcription rates throughout prophase I. As zygotene cells transitioned into pachytene, we observed a ∼4-fold increase in uridine signal intensity per cell, reflecting a marked rise in RNA synthesis. Although transcription decreased in diplotene cells, the reduction did not reach statistical significance. Cells in diakinesis, anaphase, and telophase of meiosis I exhibited negligible 5-EU incorporation, consistent with transcriptional silencing during these stages (*30*) and therefore served as negative controls relative to LZ and PD cells.

Examination of pyrimidine nucleotide pool levels (**Fig. 6D**) in LZ versus PD cells revealed a pronounced decrease in the PD population, indicating markedly increased nucleotide utilization. Together, these findings support a developmentally enforced sequence in which germ cells synthesize nucleotides during the LZ stage and, upon differentiation into PD cells, suppress nucleotide biosynthesis and repurpose these pools to meet escalating transcriptional demands.

### *In vivo* inhibition of pyrimidine synthesis prevents progression from zygotene to pachytene during first-wave spermatogenesis

We next aimed to examine how inhibiting pyrimidine biosynthesis affects male meiosis *in vivo*. Because the BTB restricts access of metabolic inhibitors to meiotic cells, we performed these experiments in prepubertal mice, during the developmental window in which the BTB is not yet established (*31*, *32*). We administered the DHODH inhibitor BRQ from postnatal day 13 to 18, during which leptotene cells differentiate into zygotene and pachytene cells (*10*) **(Fig 7A).**

**Fig. 7.**
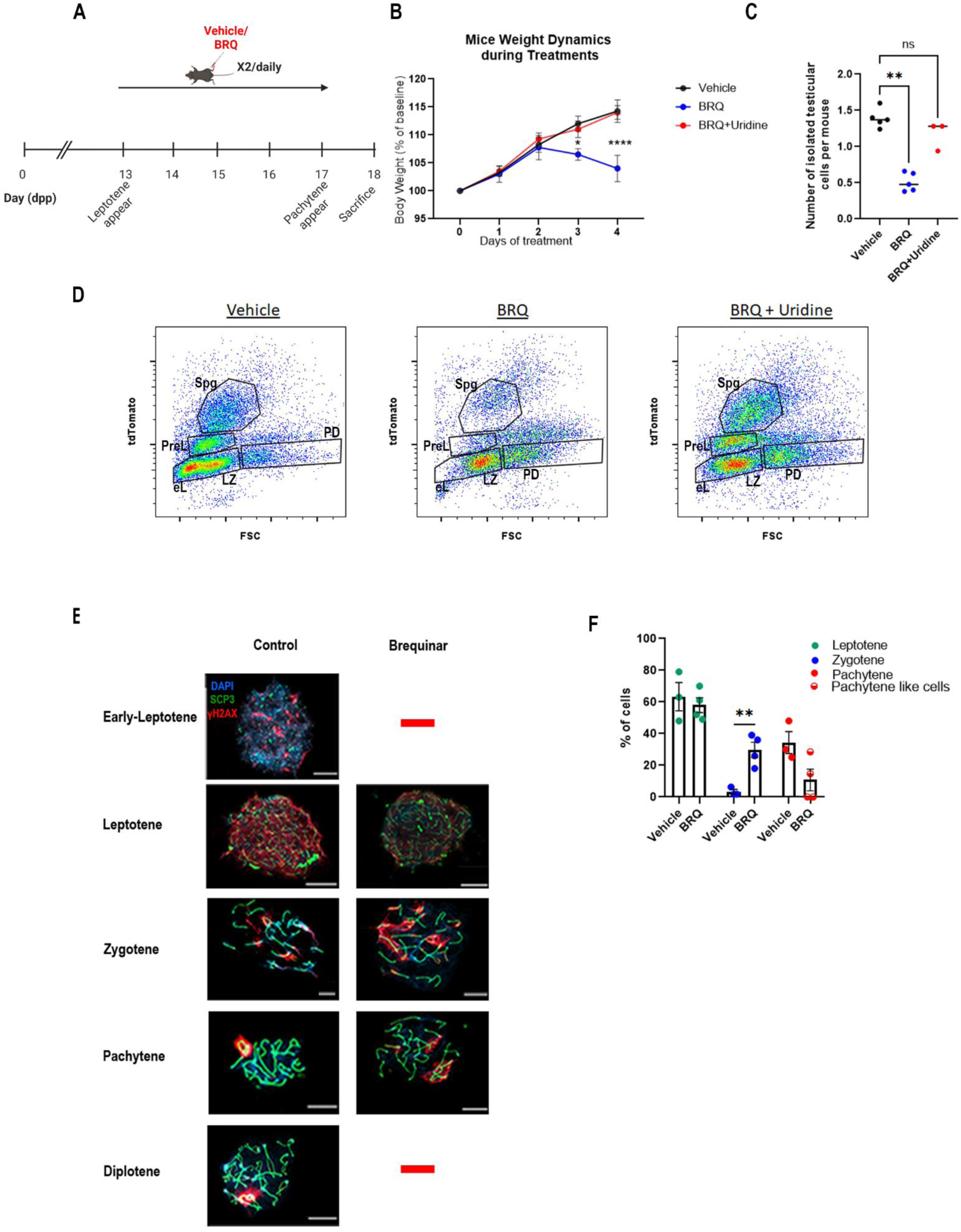
Inhibition of pyrimidine synthesis *in vivo* prevents progression from the zygotene to pachytene stage. **(A)** Experimental scheme for *in vivo* inhibition of pyrimidine biosynthesis. Prepubertal mice were treated twice daily with vehicle or BRQ **±** uridine supplementation from postnatal day (P)13-P18, spanning the developmental window in which leptotene cells emerge and progress through zygotene toward pachytene, prior to full establishment of the BTB. This figure was created with BioRender.com. (B) Mouse weight dynamics during treatment (normalized to baseline, 100%). BRQ-treated mice show reduced weight gain by day 4 of treatment, which is rescued by co-administration of uridine (Student’s T-test). **(C)** Total number of cells per mouse isolated from testes at the treatment endpoint (P18). BRQ reduces testis cell yield, and uridine co-treatment restores cell yield toward vehicle levels ANOVA. **(D)** Representative flow-cytometry profiles of germ cells isolated from Stra8-tdTomato testes following vehicle, BRQ, or BRQ+uridine treatment. Gates denote Spg, PreL, early leptotene (eL) combined with LZ and PD populations. (1 out of 3 representative experiments). **(E)** Representative nuclear spreads from sorted populations stained for SCP3 (synaptonemal complex; green), γH2AX (DSBs/sex body; red), and DNA (DAPI; blue) across meiotic stages. In vehicle-treated mice, pachytene nuclei show fully synapsed SCP3 tracks with γH2AX confined to the sex body. In BRQ-treated mice, cells within the PD gate display aberrant pachytene-like morphology with dispersed γH2AX staining. Early leptotene and diplotene cells were not detected in BRQ treated group. Scale bars=10 µm. **(F)** Quantification of meiotic-stage composition among analyzed spreads, Bars show mean ± SEM (n =3-4 independent experiments) (Student’s T-test). **P < 0.01; ****P < 0.0001; n.s., not significant.

To address potential drug toxicity, we monitored pup growth and observed a significant reduction in weight gain by day 17 in BRQ-treated mice (**Fig. 7B**). Co-administration of uridine fully rescued this phenotype, indicating that the growth defect reflects nucleotide depletion rather than broader toxic effects. Similarly, the total number of cells isolated from testes on day 18 was reduced by ∼2.8-fold in BRQ-treated mice, an effect completely reversed by uridine supplementation (**Fig. 7C**).

FACS analysis of testes from Stra8-tdTomato mice, in which germ cells are fluorescently labeled, revealed that BRQ treatment substantially altered germ cells’ composition (**Fig. 7D**). Specifically, BRQ caused a marked reduction in Spg, PreL, and early leptotene (eL) cells, whereas the LZ population was comparatively less affected.

To further characterize these changes, LZ and PD cells were subjected to nuclear spread analysis and immunostaining for the synaptonemal complex protein SCP3 and γH2AX. Leptotene cells were identified by dispersed γH2AX staining, reflecting physiologic double-strand breaks (DSBs), whereas pachytene cells displayed fully synapsed SCP3-labeled chromosomes and γH2AX restricted to the sex body (**Fig. 7E**).

In BRQ-treated mice, cells within the LZ gate consisted of morphologically normal leptotene and zygotene cells. The PD gate contained abnormal pachytene-like cells with dispersed γH2AX staining, in contrast to vehicle-treated mice, which showed typical pachytene and diplotene cells with γH2AX confined to the sex body. The significant enrichment of zygotene cells in the BRQ-treated group (**Fig. 7F**), together with the appearance of defective pachytene-like cells, and lack of diplotene cells indicates that BRQ treatment disrupts meiotic progression during the pachytene stage. Notably, mice treated with BRQ and uridine displayed FACS distributions and morphologies nearly indistinguishable from vehicle controls (**Fig. 7D, S7**).

Together, these findings demonstrate that inhibition of pyrimidine synthesis not only reduces the pool of replicative cells-including Spg and preL-but also blocks meiotic progression beyond the zygotene stage.

### Pachytene cells rely exclusively on nucleotide pools generated during earlier developmental stages rather than on external sources

A key question arising from our findings is whether the pyrimidine (and potentially purine) nucleotides required for proper differentiation of pachytene cells are supplied exclusively by earlier meiotic stages or whether they can also be supplemented from external sources, such as Sertoli cells. Our transcriptomic analysis revealed that among plasma-membrane nucleoside transporters, ENT1 (encoded by SLC29a1) is the only one robustly expressed across germ-cell populations-from differentiating spermatogonia through round spermatids (**Fig.8A**)-suggesting the possibility of nucleoside import from the somatic environment. Of note SLC29a2 that codes for ENT2 was only negligibly expressed.

**Fig. 8.**
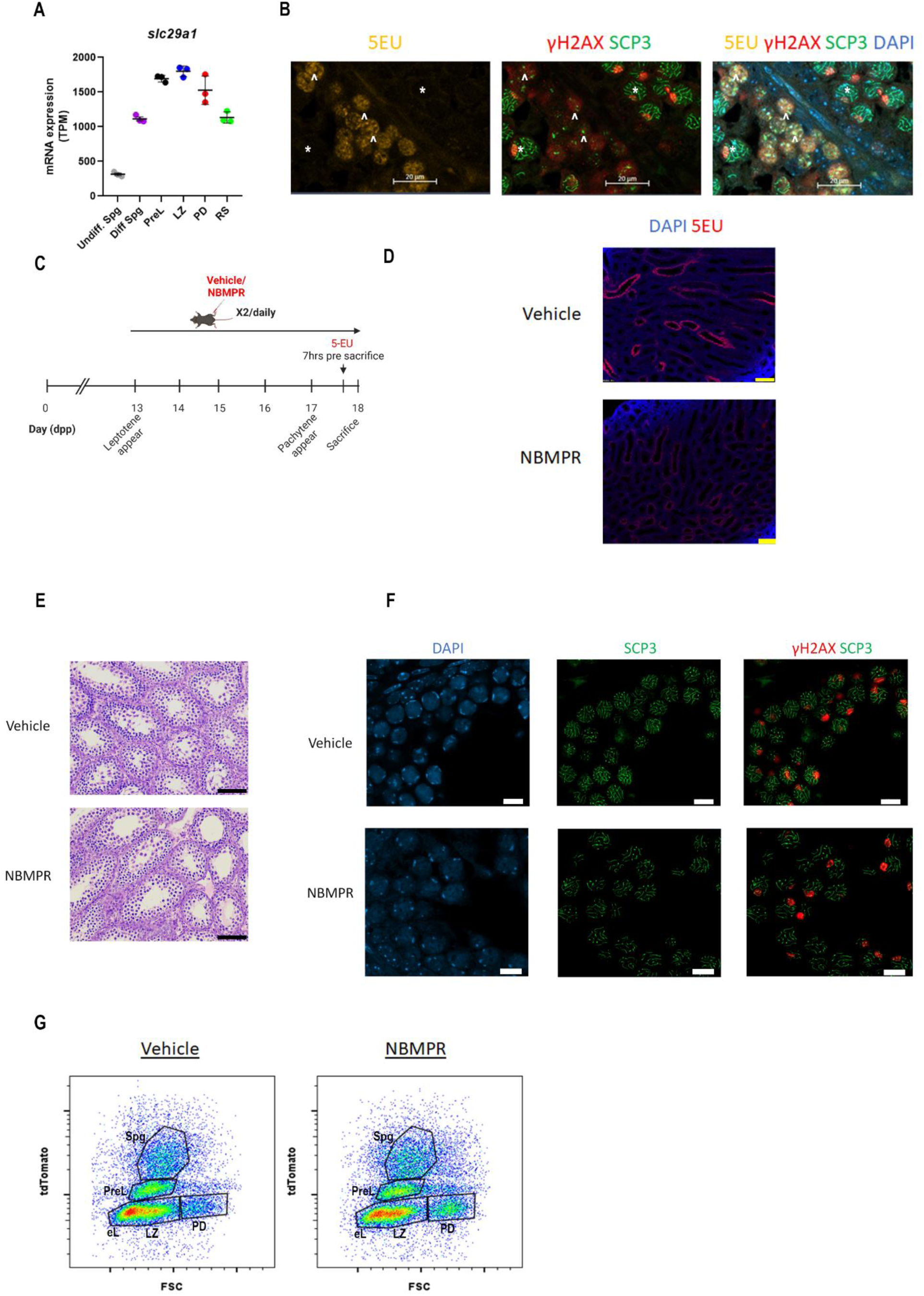
Pachytene cells rely on nucleotide pools generated during earlier developmental stages rather than on external nucleoside sources. **(A)** Expression of the nucleoside transporter ENT1 (Slc29a1) across germ-cell populations, showing Slc29a1 mRNA abundance in LZ and PD cells (RPKM). **(B)** Representative images of seminiferous tubules from P18 mice after 5-EU injection showing 5-EU signal (yellow), meiotic markers γH2AX (red) and SCP3 (green), and merged channels including DAPI (blue). 5-EU incorporation is detected in leptotene (arrowheads, ^) but is absent from pachytene cells (asterisks, *). Scale bars = 20 µm. **(C)** Scheme for pharmacologic inhibition of nucleoside uptake. Prepubertal mice were treated twice daily with vehicle or the selective ENT1 inhibitor NBMPR from P13-P18; mice received a 5-EU pulse prior to sacrifice (timing as indicated) to assess nucleoside uptake *in vivo*. This figure was created with BioRender.com. **(D)** Representative paraffin sections showing 5-EU incorporation (red) and DNA (DAPI, blue) in testes from vehicle- and NBMPR-treated mice. NBMPR markedly reduces 5-EU labeling, confirming effective inhibition of nucleoside uptake. Scale bar=100 µm. **(E)** Hematoxylin and eosin staining of testis sections from vehicle- and NBMPR-treated mice, showing comparable seminiferous tubule organization and no overt disruption of meiotic progression. Scale bar=100 µm. **(F)** immunofluorescence staining for SCP3 (green), γH2AX (red), and DAPI (blue) of vehicle- and NBMPR-treated testes. Pachytene cells display normal morphology with γH2AX confinement to the sex body in both conditions, indicating preserved meiotic progression despite blocked nucleoside uptake. Scale bar=10 µm. **(G)** Representative flow-cytometry profiles of germ cells from Stra8-tdTomato testes following vehicle or NBMPR treatment. Gates denote Spg, PreL, eL combined with LZ and PD populations; profiles are indistinguishable between conditions.

To determine which germ-cell populations are capable of nucleoside uptake, we injected 18-day-old mice with 5-EU and monitored incorporation into RNA using click chemistry. 5-EU incorporation was detectable only in germ cells up to the leptotene stage, with no detectable signal in pachytene cells (**Fig. 8B**).

We next tested whether nucleoside uptake through ENT1 is required for meiosis. Pups were treated from P13 to P18 with the selective ENT1 inhibitor nitrobenzylthioinosine (NBMPR) (*12*, *33*) (**Fig. 8C**). NBMPR treatment caused a pronounced reduction in 5-EU incorporation into germ cells, as observed in paraffin-embedded testis sections (**Fig. 8D**), confirming the efficacy of ENT1 inhibition. Despite completely blocking 5-EU uptake, NBMPR treatment did not impair progression through meiosis I. Hematoxylin and eosin staining of testis sections appeared normal **(Fig. 8E**), and pachytene cells in both NBMPR- and vehicle-treated mice displayed typical morphology with γH2AX confined to the sex body (**Fig. 8F**). Moreover, the FACS profiles of NBMPR-treated and vehicle-treated testes were indistinguishable (**Fig. 8G**).

Together, these findings indicate that nucleotides synthesized *de novo* during the early stages of meiotic prophase I are sufficient to support normal germ-cell differentiation. External sources of nucleosides-such as Sertoli-cell-derived pools-do not make a significant contribution to sustaining nucleotide levels in pachytene cells.

### Cross-species transcriptomics analysis reveals an evolutionarily conserved shutdown of nucleotide biosynthesis during meiosis

Our results demonstrated that PD cells are unable to synthesize pyrimidine nucleotides in mice. To elucidate the molecular basis underlying this metabolic limitation, we examined the transcriptional profiles of rate-limiting enzymes involved in both purine and pyrimidine nucleotide biosynthesis using our bulk RNA-seq data (GSE162740) from LZ and PD populations **(Fig. 9A-B)**. This analysis revealed a marked and significant reduction in the expression of key enzymes required for purine synthesis, including *Ppat*, *Gart*, and *Mthfd1*, as well as pyrimidine synthesis enzymes *Cad*, *Dhodh*, *Umps*, and *Ctps1*, during the transition from LZ to PD.

**Fig. 9.**
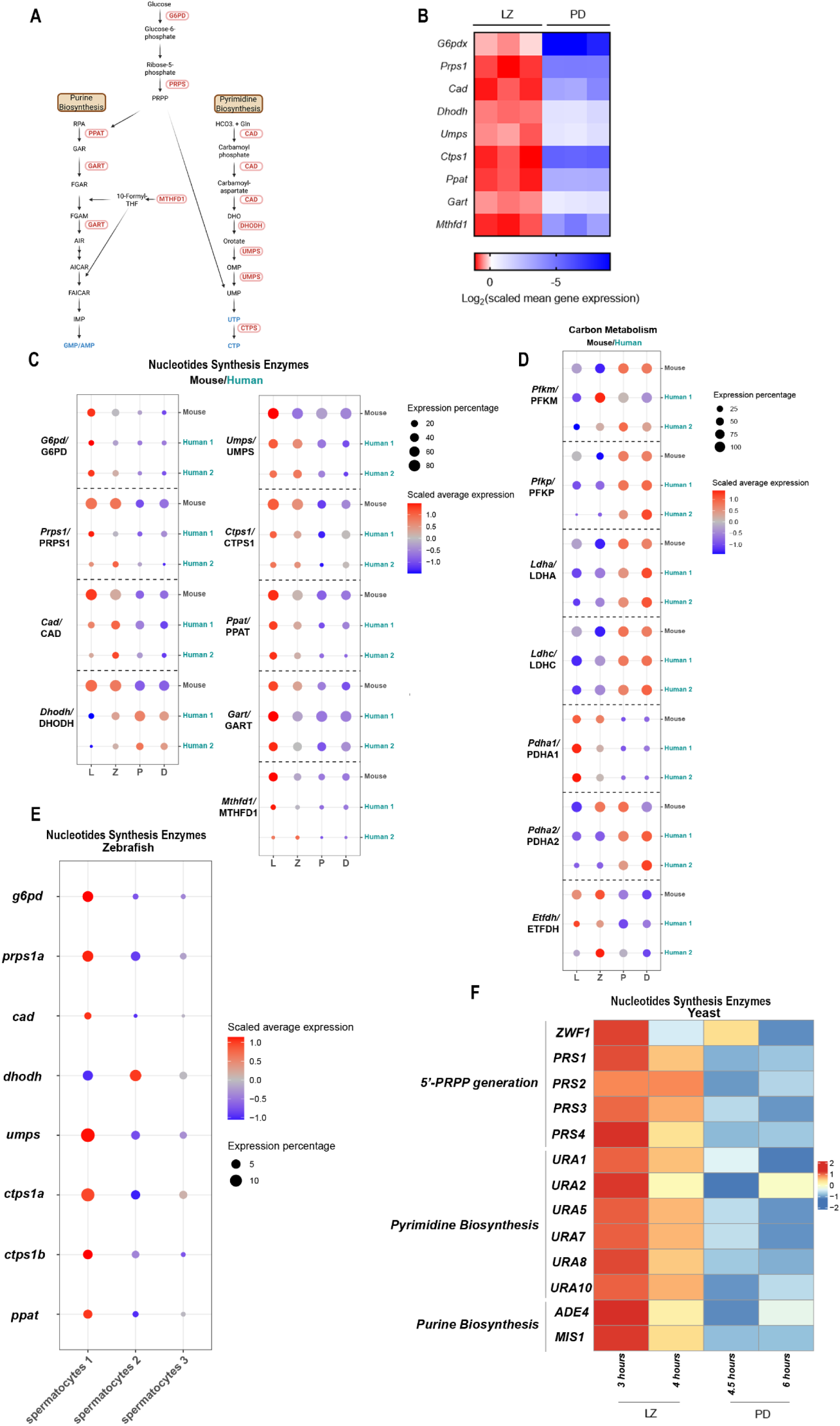
Cross-species transcriptomics reveals an evolutionarily conserved shutdown of nucleotide biosynthesis during late meiotic prophase I. **(A)** Scheme of *de novo* purine and pyrimidine nucleotide biosynthesis highlighting key rate-limiting enzymes analyzed in this study. This figure was created with BioRender.com. **(B)** Heat map of bulk RNA-seq expression (GSE162740) of nucleotide biosynthetic genes in LZ and PD populations, showing coordinated transcriptional repression of enzymes required for purine synthesis (*Ppat*, *Gart*, *Mthfd1*) and pyrimidine synthesis (*Cad*, *Dhodh*, *Umps*, *Ctps1*) during the LZ-to-PD transition, together with suppression of *G6pd* and *Prps1* in mice. **(C)** Cross-species single-cell RNA-seq meta-analysis of nucleotide biosynthesis genes in mouse (GSE148032) and human (two data sets: GSE112013 and GSE106487) prophase I subpopulations. Dot plots show scaled average expression (color) and fraction of expressing cells (dot size) across annotated stages. **(D)** Cross-species expression dynamics of carbon-metabolism genes associated with the LZ-to-PD metabolic switch. **(E)** Single-cell RNA-seq analysis of zebrafish (GSE275361) showing downregulation of genes involved in 5-PRPP generation (*g6pd*, *prps1a*, *prps1b*), pyrimidine biosynthesis (*cad, dhodh, umps, ctps1a, ctps1b*), and purine biosynthesis (*ppat*) upon transition from early to late prophase I (LZ to PD, based on marker-defined populations). **(F)** Bulk mRNA-seq time course of synchronous meiosis in *S. cerevisiae* (GSE34082). Heat map shows repression of genes involved in 5-PRPP generation (*ZWF1, PRS1-4),* pyrimidine biosynthesis (*URA1/2/5/7/8/10*), and purine biosynthesis (*ADE4*, *MIS1*) during progression into late meiotic prophase.

In addition, the X-linked metabolic enzymes *G6pd* and *Prps1*, which are essential for the generation of 5-phosphoribosyl-1-pyrophosphate (5-PRPP), exhibited pronounced transcriptional silencing at the PD stage. This repression is consistent with MSCI. Collectively, these findings indicate that a coordinated transcriptional shutdown of both purine and pyrimidine biosynthetic pathways underlies the inability of PD cells to autonomously synthesize nucleotides.

Meiosis is widely regarded as a conserved and fundamental feature of eukaryotic sexual reproduction, with core regulatory and cellular processes preserved across evolutionarily distant lineages (*34*, *35*). We therefore sought to determine whether the shutdown of nucleotide biosynthesis observed in PD cells represents a conserved feature of meiosis across species. To address this, we performed a cross-species meta-analysis of three previously published single-cell RNA-seq datasets-one mouse (*36*) (GSE148032) and two humans (*37*, *38*) (GSE112013 and GSE106487)-each with clear annotation of prophase I subpopulations.

Consistent with our bulk RNA-seq findings, single-cell analyses of mouse data revealed a downregulation of nucleotide biosynthetic enzymes during the LZ-to-PD transition **(Fig. 9C**, subpopulations markers depicted in **S9A).** Expression of *Ppat*, *Gart*, and *Mthfd1* peaked during the LZ stage and was markedly suppressed upon entry into PD. Similarly, *Cad*, *Umps*, and Ctps1 displayed robust expression in early prophase I followed by pronounced downregulation in pachytene and diplotene cells. Both human studies showed a very similar pattern of changes. The only species-specific behavior could be noted in *DHODH*/Dhodh expression. It increased during the LZ-to-PD transition in humans while decreased in mouse. Notably, the X-linked genes *G6PD*/G6pd and *PRPS1*/Prps1 were consistently silenced following LZ progression in both organisms, reinforcing the conserved impact of sex chromosome regulation on meiotic metabolism.

We also examined the change in the expression of enzymes that we pinpointed as involved in the metabolic switch during LZ to PD transition. In contrast to the repression of nucleotide biosynthesis, glycolytic enzymes *Pfkm/PFKM*, *Pfkp/PFKP*, *Ldha/LDHA*, and *Ldha/LDHC* transitioned from low basal expression during LZ to high transcript abundance in PD cells in both mouse and human. Moreover, a conserved isoform switch was observed within the PDC, characterized by silencing of the X-linked *Pdha1/PDHA1* gene alongside upregulation of the autosomal *Pdha2/PDHA2* gene during later stages of prophase I. in addition, ETFDH expression was silenced upon LZ to PD transition in both species. **(Fig. 9D)**

Based on these observations, we hypothesized that the metabolic differences between LZ and PD cells may arise from intrinsic constraints imposed on PD cells, particularly their inability to express X-linked metabolic enzymes such as *G6pd/G6PD* and *Prps1/PRPS1* because of MSCI. To test this idea, we extended our analysis to zebrafish (*Danio rerio*) and yeast (*Saccharomyces cerevisiae*), eukaryotes that lack sex chromosomes and do not undergo MSCI.

Analysis of previously published single-cell RNA-seq data (GSE275361) from zebrafish (36) showed that genes involved in 5-PRPP generation (*g6pd, prps1a,* and *prps1b*), pyrimidine synthesis (*cad, umps, ctps1a,* and *ctps1b*), and purine synthesis (*ppat*) were significantly downregulated during the transition from early to late prophase I (**Fig. 9E**), corresponding to the LZ-to-PD transition as defined by the relevant markers (**Fig. S9B**). These results were similar to above mouse and human data. *gart*, *mthfd1a* and *mthfd1b* expression was negligible in zebrafish cells.

To test whether repression of nucleotide biosynthesis occurs in a distantly related eukaryotic unicellular organism, we analyzed previously published mRNA-seq bulk data (GSE 34082) from *S. cerevisiae* (*39*). In this system, synchronous meiosis was induced by incubating cells in SPO medium to induce sporulation. Entry into pachytene stage was accompanied by significant downregulation of genes involved in 5’-PRPP generation (ZWF1 and *PRPS1-4*), pyrimidine biosynthesis (*URA1, URA2*, *URA5, URA7, URA8, URA10*), purine biosynthesis (*ADE4, MIS1*) (**Fig. 9F, S9C**). The presence of a comparable transcriptional shutdown in zebrafish and yeast, which both lack sex chromosomes and does not undergo MSCI, demonstrates that repression of nucleotide biosynthetic capacity during late meiotic prophase is evolutionarily conserved and cannot be attributed to MSCI.

Together, these results demonstrate that the suppression of nucleotide biosynthesis during late prophase I is an ancient evolutionary conserved feature of meiosis, across diverse eukaryotes, operating independently of sex chromosome presence. While MSCI reinforces this repression in mammals through silencing of X-linked metabolic genes, analogous transcriptional downregulation in zebrafish and yeast indicates that this metabolic transition reflects a fundamental constraint or regulatory principle intrinsic to meiotic progression.

## Discussion

Using our recently published method for isolating spermatogenic cells at defined developmental stages, we were able to characterize metabolic changes across spermatogenesis. Earlier metabolic studies were largely constrained by the cell types that could be isolated with available methodologies. Nevertheless, these works established a key principle: lactate supplied by Sertoli cells serves as an important fuel for pachytene spermatocytes (*40*) and round spermatids (*41*, *42*), as reviewed in Boussouar and Benahmed (*43*). Our findings are consistent with this model, and further show that cells preceding pachytene-specifically the LZ population-are unable to effectively utilize lactate and instead rely on alternative fuels, including glutamine and fatty acids.

Whitmor et. al attempted to overcome stage-availability limitations by profiling ribosome-associated mRNAs during synchronized first-wave spermatogenesis in prepubertal mice, up to the leptotene stage, coupled with network modeling (*44*). That study focused primarily on retinoic acid metabolism, highlighting the value of higher-resolution stage-resolved approaches at early stages of spermatogenesis.

By integrating transcriptomic, proteomic, and metabolomic datasets with ^13^C-tracing, we identify additional metabolic features that accompany progression through meiosis and revealed major metabolic transitions between the LZ and PD populations. These transitions involve not only shifts in fuel preference but also changes in mitochondrial abundance and composition, consistent with our observed differences in oxygen consumption rate (OCR). In line with this, mitochondrial morphological remodeling during spermatogenesis has been reported previously (reviewed in (*45*). In our study, mitochondrial proteomics supports a functional reorganization that enhances amino acid and fatty-acid metabolism together with increased energy production via the TCA cycle and oxidative phosphorylation during the LZ-to-PD transition.

Analysis of key metabolic enzymes in glycolysis, the pentose phosphate pathway (PPP), and the TCA cycle yielded several notable observations. Genes encoding testis-specific isoforms of glycolytic enzymes (*Gapdhs, Pgk2*, *Ldhc*), as well as the testis-specific *Pdh* subunit *Pdha2*, are highly expressed at the PD stage. This expression pattern is consistent with increased glycolytic flux, lactate utilization, and efficient channeling of pyruvate into the TCA cycle in PD cells compared to the preceding LZ stage.

The LZ-to-PD metabolic program we describe-specifically, reduced expression of PFK and PDC components in LZ cells and suppression of the PPP in PD cells-may reflect a developmental requirement to downregulate *de novo* nucleotide synthesis at later stages and instead utilize nucleotides synthesized earlier (**Fig. 10**). Notably, PD cells also exhibit a marked shutdown of fatty-acid β-oxidation, accompanied by reduced abundance of ETFA, ETFB, and ETFDH, which are required for electron transfer and regeneration of FAD. One possible explanation is that limiting β-oxidation supports lipid anabolism at this stage, when cells undergo substantial biomass expansion and membrane synthesis.

**Fig 10.**
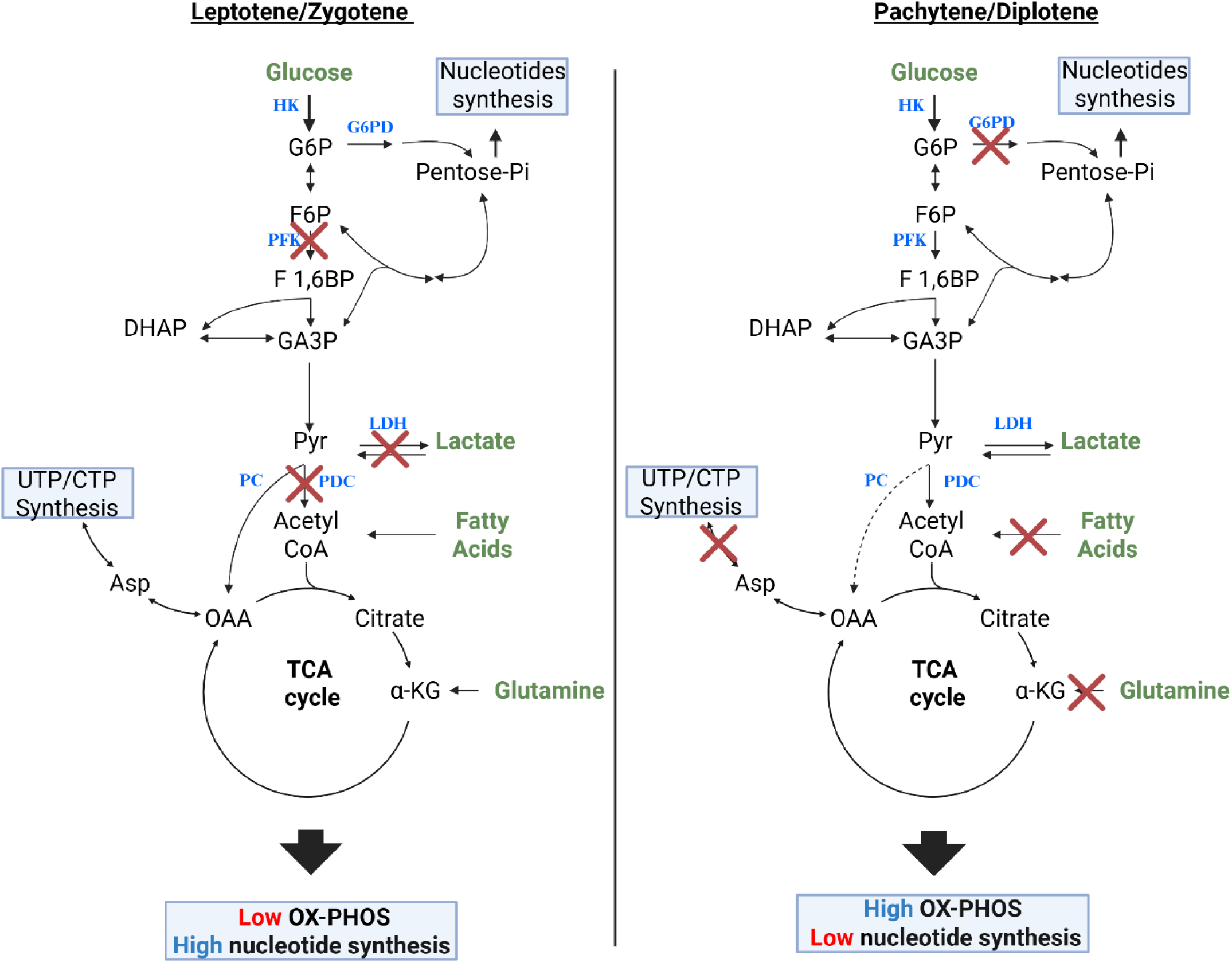
Summary model of the LZ–to-PD metabolic transition during meiotic prophase. **I** Scheme integrating multi-omics and isotope-tracing results to depict stage-specific fuel usage and biosynthetic output in LZ (left) versus PD (right) spermatocytes. Red “X” symbols indicate pathways or fuel routes that are reduced or inactive in the indicated stage; arrows indicate predominant carbon flow.

Metabolic regulation in PD-stage spermatocytes is further constrained by meiotic sex chromosome inactivation (MSCI), which transcriptionally silences the X chromosome during pachytene. In mouse, the X chromosome encodes 978 protein-coding genes (MGI_EntrezGene.rpt), and 467 carry Gene Ontology Biological Process annotations under “metabolic process” (GO:0008152), creating a potential bottleneck for pathways that depend on X-linked enzymes. Consistent with this idea, our data show that the PPP can be effectively shut down in PD cells coincident with suppression of G6PD at both the transcript and protein levels.

A well-described evolutionary solution to MSCI is retroposition or duplication of X-linked genes to autosomes, generating autosomal paralogs/retrogenes that remain expressed when the X is silent. Several such autosomal copies show testis-specific expression and are abundant at the PD stage (*46–48*). In addition to gene relocation, PD cells likely rely on multiple regulatory layers to match metabolic supply and demand during the LZ-to-PD transition. Large-scale transcriptome remodeling accompanies this transition, with on the order of ∼10,000 genes changing significantly in expression (*9*, *49*). Moreover, extensive alternative splicing-a particularly prominent feature of the testis (*50*)-together with mRNA stabilization (*51*) and other post-transcriptional mechanisms (*52*), provide substantial capacity to tune protein output and support metabolic adaptation despite constraints imposed by MSCI.

The evolutionary conservation of nucleotide biosynthesis shutdown during late meiotic prophase, observed in organisms that predate the emergence of sex chromosomes, suggests that this metabolic program did not arise as a consequence of X-chromosome regulation. Rather, meiotic silencing of nucleotide biosynthesis appears to represent an ancient, intrinsic feature of meiosis, upon which meiotic sex chromosome inactivation was later superimposed in mammals.

Finally, our comparative analysis of nucleotide biosynthesis gene expression across diverse organisms-from yeast to humans-suggests that suppression of these pathways during late stages of prophase I of both male meiosis and yeast sporulation is evolutionarily conserved. An important goal for future work will be to determine how this shutdown is regulated and to clarify its functional role during meiotic progression.

This study combines multi-omics measurements and isotope tracing in purified spermatogenic populations, but several limitations should be considered. First, metabolic states can be perturbed by tissue dissociation, FACS sorting, and short ex vivo incubations; although we minimized these effects and validated key findings across conditions, some metabolite abundances and fluxes may not fully reflect in situ physiology. Second, our stage-resolved analyses compare gate-defined LZ and PD populations, which each encompass cellular heterogeneity and a temporal continuum within prophase I; higher-resolution approaches (e.g., finer sub-staging, time-resolved sampling, or single-cell multi-omics) will be needed to pinpoint when specific metabolic switches occur. Third, our conclusions about nucleotide “storage” are inferred from pathway activity, labeling patterns, and inhibitor experiments; we do not directly define the chemical form, subcellular compartmentalization, or turnover kinetics of the nucleotide pools that support PD transcriptional demands. Fourth, our *in vivo* pharmacology was performed in prepubertal mice to bypass the fully established blood-testis barrier, and thus may not capture all regulatory constraints present in the adult testis. Finally, the cross-species conservation analysis is based on transcriptomic signatures; functional tests in non-mammalian systems will be required to establish whether repression of nucleotide biosynthesis is mechanistically conserved and to determine how it is coupled to meiotic progression.

## METHODS

### Ethics statement

All mice were bred at the Hebrew University- Hadassah Medical School (Jerusalem, Israel), and protocols for their care and use were approved by the Institutional Animal Care and Use Committee of the Hebrew University (approval #MD-22-16877-2).

### Mice

Mice were housed under a regular 12-h/12-h dark/light cycle with ad libitum access to food and water. Stra-icre ( B6.FVB-Tg(Stra8-icre)1Reb/LguJ) male mice (stock 017490) (*53*), tdTomato (B6;129S6-Gt(ROSA)26Sor^tm9(CAG-tdTomato)Hze^/J females(stock 007905) were purchased from Jackson Laboratories. All mice are on C57Bl background. Hemizygous Stra-icre males were mated with homozygous tdTomato and males which were positive for Stra-icre were used as a source of testis tissue (Stra8-tdTomato mice). Genotyping was done by RT PCR according to the suppliers’ protocol.

### Cells Isolation

Germ cells isolation was performed, as described in our previous study in details(*9*). In short, adult mice, 12-16 weeks old, or 18 days old pups were euthanized with pental, and cervical dislocation was performed. Testis were removed, put into cold Hanks’s solution and seminiferous tubules were released from the tunica albuginea. Tissue digestion was performed in two steps. First, with 4 min incubation with collagenase (C5138 Merck) 1 mg/ml and DNAse (DN-25 Merck) 0.28 mg/ml, followed by 20 min digestion in 33 deg in trypsin-EDTA (D705 Diagnovum) supplemented with 0.93 mg/ml DNAse. The resulting suspension was passed through 40 µm cell strainer, washed in medium and incubated with dead cells removal kit (130-090-101 Miltenyi) for 15 min in room temperature. Live cells were collected according to the manufacturer’s instructions and washed in PBS with 5% serum followed by dispersal in sorting buffer. Sorting buffer consisted of PBS with 25 mM HEPES, 5% FBS, 0.05 mg DNAse and 0.75 mM MgCl2.

### Flow Cytometry Sorting

After live cell isolation, cells were washed in 2% dialyzed serum in PBS, resuspended in sorting buffer at the density of 10 million cells / 1 ml. During sorting, cells were collected into 5 ml tubes, which were kept overnight in 4°C, inverted with 2ml of different collecting buffers. Collecting buffer for the immediate extraction of RNA consisted of PBS with 2% albumin. Collecting buffer for the immediate extraction of metabolites and LC-MS analysis consisted of PBS with 5% dialyzed FBS. For overnight recovery from sorting stress, cells were collected in culture medium consisting of MEM Eagle medium supplemented with glutamine 2 mM, lactate 5 mM, pyruvate 5 mM, HEPES 15 mM, dialyzed serum 5% and antibiotic-antimycotic solution (Thermo Scientific, 15240096).

Cells were sorted using a BD FACSAria™ III sorter, as described previously (*9*). Debris exclusion was done based on light scattering. Tomato intensity (yellow-green laser, bandpass 582/15) of cells versus forward scatter (FSC) was used to differentiate between the different populations, **(Fig S1)**. Doublet’s exclusion in each population was performed on the basis of FSC width/area (FWA), followed by side scatter (SSC) width/area (SWA) and tomato width/area (PWA).

Four-way sorting was performed using 100-micron nozzle and threshold was kept in the range of 2500-3000 events/second. PD cells were fragile and sensitive to high pressures that can be developed during sorting; thus, the above conditions were found to be appropriate to keep the PD cells alive.

### Metabolomic changes along spermatogenesis

Metabolomic changes along spermatogenesis were studied in two setups: in freshly isolated cells and in cells after over-night recovery in MEM Eagle (Sartorius 01-025) supplemented with 2 mM each glutamine, L-lactate and pyruvate and 5% dialyzed serum. This was followed by 2 h incubation in the synthetic seminiferous tubule fluid sSTF (see Table S2). Cells were washed twice in PBS and extracted with acetonitrile/methanol/water (5:3:2) and frozen in −80 deg. LC-MS measurements are described below.

### LC-MS

Over-night frozen samples were thawed, vortexed and centrifuged for 20 minutes at 4°C at 20,000Xg. Pellets were used to measure protein content. Extracted metabolites were transferred to new, −80°C precooled, tubes and chromatographically separated on a SeQuant ZIC-pHILIC column (2.1 × 150 mm, 5 μm bead size, Merck Millipore). Flow rate was set to 0.2 mL/min, column compartment was set to 30°C, and autosampler tray was maintained at 4°C. Mobile phase A consisted of 20 mM ammonium carbonate with 0.01% (v/v) ammonium hydroxide. Mobile Phase B was 100% acetonitrile. The mobile phase linear gradient (%B) was as follows: 0 min 80%, 15 min 20%, 15.1 min 80%, and 23 min 80%. A mobile phase was introduced to Thermo Q-Exactive mass spectrometer with an electrospray ionization source working in polarity switching mode. Metabolites were analyzed using full-scan method in the range 70-1,000 m/z and with a resolution of 70,000. Positions of metabolites in the chromatogram were identified by corresponding pure chemical standards. Data were analyzed using the MAVEN software suite. Relative metabolite abundances were quantified from peak area and normalized to cell volumes. Principal component analysis and heatmaps were generated using Metaboanalyst software (*54*, *55*).

### Oxygen consumption rate (OCR) assessment

OCR was measured using a Seahorse XFe96 Analyzer system.

Cells were enzymatically isolated as described above and collected into PBS with 5% Fetal bovine serum (not dialyzed). Cells were washed twice in PBS, then adhered to the cell plate covered with Cell-Tak (354240 Corning). Cell-Tak coating of the cell plate was as following: 9µl of Cell-Tak (1.16mg/ml) was introduced into 325 µl of 0.1M sodium bicarbonate, pH was corrected with 4.5 µl NAOH 1N. 10 µl of this solution was put on 96 wells plate and incubated for 20 min in rt, followed by two washes with 200 µl of water and drying.

Cells were seeded in 30µl salt solution consisting of 142 mM NaCl, 5.4 mM KCl, 0.91 mM NaH2PO4, 1.8 mM CaCl2, 0.8 mM MgSO4 and HEPES 5mM, pH 7.35. PD cells are sensitive to Mg concentration. Higher than 0.8 mM Mg can cause high death rate. The amount seeded cells was: Spg 120 000, LZ 200 000, PD 160 000 and RS 180 000 cells/well.

Cells were centrifuged twice for 1 min at 200g, acceleration 5, no brake, each time at a different direction, followed by incubation for 30 min in 37°C in incubator without CO_2_. After the incubation, OCR measurement buffer was added stepwise: 50 µl and 100 µl in order to prevent cells detachment. OCR measurement buffer consisted of 142 mM NaCl, 5.4 mM KCl, 0.91 mM NaH2PO4, 1.8 mM CaCl2, 0.8 mM MgSO4 and HEPES 5mM, 0.4% albumin FA free, 2mM l-carnitine hydrochloride, pH 7.35

After cartridge calibration, basal OCR measurements were performed in three cycles of 30 sec mix, 3 min wait and 3 min measurement. Injection ports were loaded with 20 µl solutions and total volume in each well was 200 µl. The concentration of fuels, uncoupler CCCP (HY-100941 MCE) and etomoxir (an inhibitor of carnitine acyl transferase 1) (HY-50202 MCE) is indicated in the legends.

### Mitochondrial content changes along spermatogenesis

Enzymatically isolated cells after treatment with dead cell removal kit were incubated for 20 min with MitoTracker Green FM (M46750 Thermo Fisher Scientific) in sorting buffer using dilution 1:33333 and analyzed in BD FACS Aria.

### Measurement of fatty acid oxidation

LZ and PD cells were sorted and regenerated over-night by incubation in MEM Eagle, as described above in the “ Metabolomic changes along spermatogenesis” section. Next, 1.5 million cells were transferred into Eppendorf tubes containing 100 µl sSTF (see **Table S2**) supplemented with 0.3 % FA-free albumin, 1mM L-carnitine, 0.2 mM palmitate and 0.66 µCi 1-^14^C-palmitate (NEC075H050UC, Revvity) and incubated for 2 h in 32 degrees. ^14^CO_2_ was trapped on Whatman filter soaked with NaOH as described in (*56*) and measured in beta counter (Beckman).

### Stable isotope tracing

Cells were sorted and recovered over-night in MEM Eagle medium, as described above in the “*Metabolomic changes along spermatogenesis*” section. Following these 1 to 1.5 million cells were transferred into Eppendorf tubes containing 200 µl sSTF (see **table S2**) supplemented with either 5 mM U-^13^C-glucose (CLM-1396-MPT-PK, Cambridge Isotope Laboratories) or 2 mM U-^13^C-glutamine (CLM-1822-H-PK, Cambridge Isotope Laboratories) and incubated for 2 h in 32 degrees.

At the end of the incubation cells were washed twice in PBS, and extracted and analyzed as above in **“***Metabolomic changes along spermatogenesis”* section.

### Estimation of the transcriptional activity of germ cells

Freshly isolated testicular cells from adult mice were prepared as described above. Cells were resuspended and allowed to recover for 2 h in Minimum Essential Medium Eagle (MEM Eagle; Sartorius, 01-025) supplemented with 2 mM each of glutamine, L-lactate, and pyruvate, and 5% serum. After recovery, 5-ethynyluridine (5-EU) was added from a 100 mM stock solution in DMSO to a final concentration of 1 mM, and cells were incubated for 2 h at 32°C.

Following 5-EU labeling, cells were washed once with PBS supplemented with 1% serum and cytospun onto glass slides. Cells were fixed for 15 min in 4% paraformaldehyde (PFA), washed once with PBS, and permeabilized for 5 min with 0.5% Triton X-100 in PBS. Incorporated 5-EU was detected using the Click-iT RNA Imaging Kit (ThermoFisher C10330) according to the manufacturer’s instructions.

After the Click-iT reaction, slides were washed once with PBS containing 1% BSA and blocked for 1 h in PBS supplemented with 3% BSA, 5% donkey serum, and 0.1% Triton X-100. Mouse anti-SCP3 (Abcam, ab97672; 1:500) was diluted in blocking buffer and applied over-night at 4°C. The next day, slides were washed three times for 5 min each with PBS containing 0.1% Triton X-100 and 0.5% BSA, followed by one wash with PBS containing 1% BSA. Alexa Fluor 488-conjugated anti-mouse secondary antibody (Abcam, ab150117; 1:200) was applied in PBS containing 1% BSA for 2 h at room temperature in the dark. Slides were washed three times for 5 min each with PBS and mounted in mounting medium containing Hoechst (1:250). Stained cells were imaged on a Zeiss LSM 980 confocal microscope using a 40× objective.

### Inhibition of pyrimidine synthesis *in vivo* in prepuberty mice

Prepubertal Stra8-tdTomato mice were divided into three groups. The control group received vehicle injections. The brequinar (BRQ) group received BRQ (Cayman, 36183) dissolved in 30% PEG-400 in PBS, administered twice daily at 11 mg/kg intraperitoneally. The BRQ+uridine group received BRQ as above plus uridine (Merck, U3003) at 500 mg/kg dissolved in PBS. Injections began on the evening of postnatal day 13 (P13) and continued through P17 (inclusive). Mice were euthanized on P18, and testis cells were isolated and sorted as described above in the “Cell isolation” and “Flow cytometry” sections. Cells from the LZ and PD gates were collected, and nuclear spreads were prepared as described previously (*9*). Spreads were stained with anti-γH2AX (1:1000; Cell Signaling, 20E39718) and anti-SCP3 (1:100; Abcam, ab97672), and imaged on a Zeiss LSM 980 Airyscan confocal microscope using a 63× objective.

### Inhibition of nucleosides uptake *in vivo* in prepuberty mice

Prepubertal Stra8-tdTomato mice were divided into two groups. The control group received vehicle injections, and the ENT1-inhibitor group received NBMPR (Merck, N2255) twice daily at 14 mg/kg intraperitoneally. NBMPR was first dissolved in DMSO and then diluted into 50% PEG-400. Injections began on the evening of postnatal day 13 (P13) and continued through P17 (inclusive). Mice were euthanized on P18, and testis cells were isolated and sorted for FCS analysis as described above in the “Cell isolation” and “Flow cytometry” sections.

In additional experiments, vehicle treated and NBMPR treated mice were injected on P18 with 5-ethynyl uridine (5-EU; 2 mg per mouse) and euthanized 7 h later. Testes were fixed in 4% formalin and embedded in paraffin. Sections were deparaffinized, subjected to antigen retrieval in citrate buffer, and permeabilized in 0.5% Triton X-100. The Click-iT reaction was performed using the ThermoFisher kit (C10330) according to the manufacturer’s instructions. Following the Click-iT reaction, sections were immunostained for γH2AX and SCP3 as described in the preceding section. Slides were scanned on an Olympus SLIDEVIEW VS200 to assess 5-EU incorporation and imaged on a Zeiss LSM 980 Airyscan confocal microscope using a 63× objective to identify specific cell populations incorporating 5-EU. Selected testes were also stained with hematoxylin and eosin (H&E) for morphological evaluation.

### Statistical Analysis

All data are presented as mean ± SEM unless otherwise indicated in the figure legends. Statistical analyses were performed using GraphPad Prism (version 10.0.0). Significance was determined as specified in the corresponding figure legends. Statistical significance was defined as *P < 0.05; **P < 0.01; ***P < 0.001; \*\*\**P < 0.0001; ns., not significant*.

### Bioinformatics-Bulk and single cells RNAseq

Processed datasets listed as SMARTdb (listed below) were obtained via SMARTdb (*57*), an integrative database for single-cell multi-omics data related to reproduction. These datasets were obtained as h5ad files and converted to Seurat objects using R (v4.4.3) with the package sceasy (v0.0.7).

For single-cell datasets, normalized expression values from the RNA assay, as provided by SMARTdb or by Sposato et al.(*58*), were used for plotting without additional transformation. In yeast, RPKM values were log-transformed with the addition of a pseudocount of one, and timepoints I-O (3-6 h) were selected based on stage-specific marker expression.

Dot plots were created with the Seurat package (v5.2.1) DotPlot function, and subsequently replotted with ggplot2 (v3.5.1) for customized presentation.

Heatmaps were visualized using the heatmap function from the ComplexHeatmap (v2.22.0) package.

Datasets used in this study.

| Species | Dataset<br>(Condition) | Reference<br>(Year) | Source | Notes |
| --- | --- | --- | --- | --- |
| Human | Normal adult testis | Guo et al., *Cell Res* (2018)(37) | SMARTdb | — |
| Human | Adult testis (healthy, obstructive azoospermia) | Wang et al., *Cell Stem Cell* (2018)(38) | SMARTdb | L1-L3 reannotated as leptotene (L) |
| Mouse | Adult testis | Zhao et al., *Nat. Commun.* (2021)(36) | SMARTdb | Fine-grained subtype annotation |
| Zebrafish | Testis developmental atlas (5-22 months) | Sposato et al., *Development* (2024)(58) | GitHub: asposato /aging_zebrafish _testis | — |
| Yeast | Meiotic time course | Brar et al., *Science* (2012)(39) | GEO: GSE34082 | Time points I-O (3-6 h) |

Table footnote:
*Processed datasets listed as SMARTdb were obtained directly from the authors upon request, as direct download of processed data objects is not available through the SMARTdb interface. Sposato et al. repository is available at https://github.com/asposato/aging_zebrafish_testis*

## ACKNOWLEDGMENTS

We thank Dr. Michael Klutstein and Dr. Michael Berger for helpful discussions and insightful comments on the manuscript. We thank Mohammad Jumaa for excellent histological assistance, Dr. Zakhariya Manevitch for microscopy analyses, and Dr. Hadas Segev-Yekutiel for FACS analysis.

## AUTHOR CONTRIBUTIONS

Conceptualization: NM, EP, Data curation: GBP, NM. Formal analysis: GBP, NM. Investigation: GBP, NM, YM. Methodology: GBP, NM, YM, EM, BS, NS, TS. Validation: GBP, NM, YM, BS, NS. Visualization: GBP, NM. Supervision: NM, TS, EP. Project administration: IS. Bioinformatics: SE, SR, OV. Funding acquisition: EP. Writing-original draft: GBP, NM, EP.

## DATA, CODE, AND MATERIALS AVAILABILITY

All data and code needed to evaluate and reproduce the results in the paper are present in the paper and/or the Supplementary Materials and/or Method section.

