## supplementary Figures for "Stage-specific metabolic rewiring coordinates nucleotide supply and demand during spermatogenesis"

**This PDF file includes:**

Figs. S1 to S9

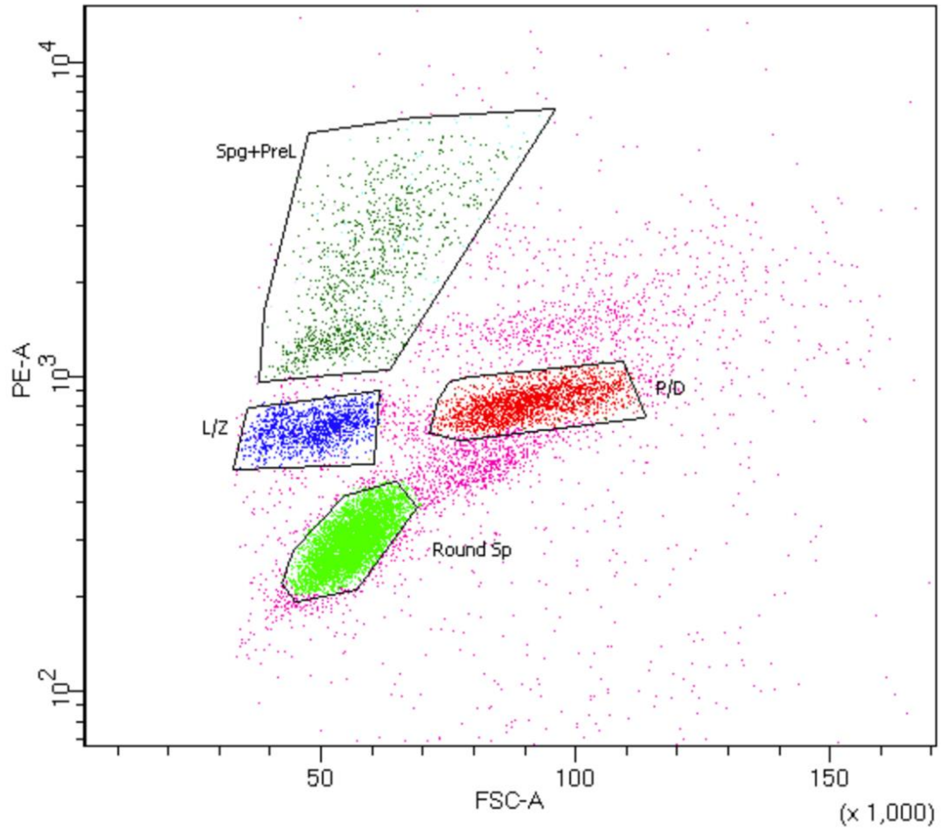


**Figure S1: Flow-cytometry isolation of spermatogenic populations for metabolomic profiling**

Representative flow-cytometry plot showing gating strategy used to isolate discrete germ-cell populations from Stra8-tdTomato mouse testes based on forward scatter (FSC-A) and PE-A (tdTomato) fluorescence. Gates correspond to spermatogonia plus pre-leptotene spermatocytes (Spg+PreL), leptotene/zygotene spermatocytes (LZ), pachytene/diplotene spermatocytes (PD), and round spermatids (Round Sp).

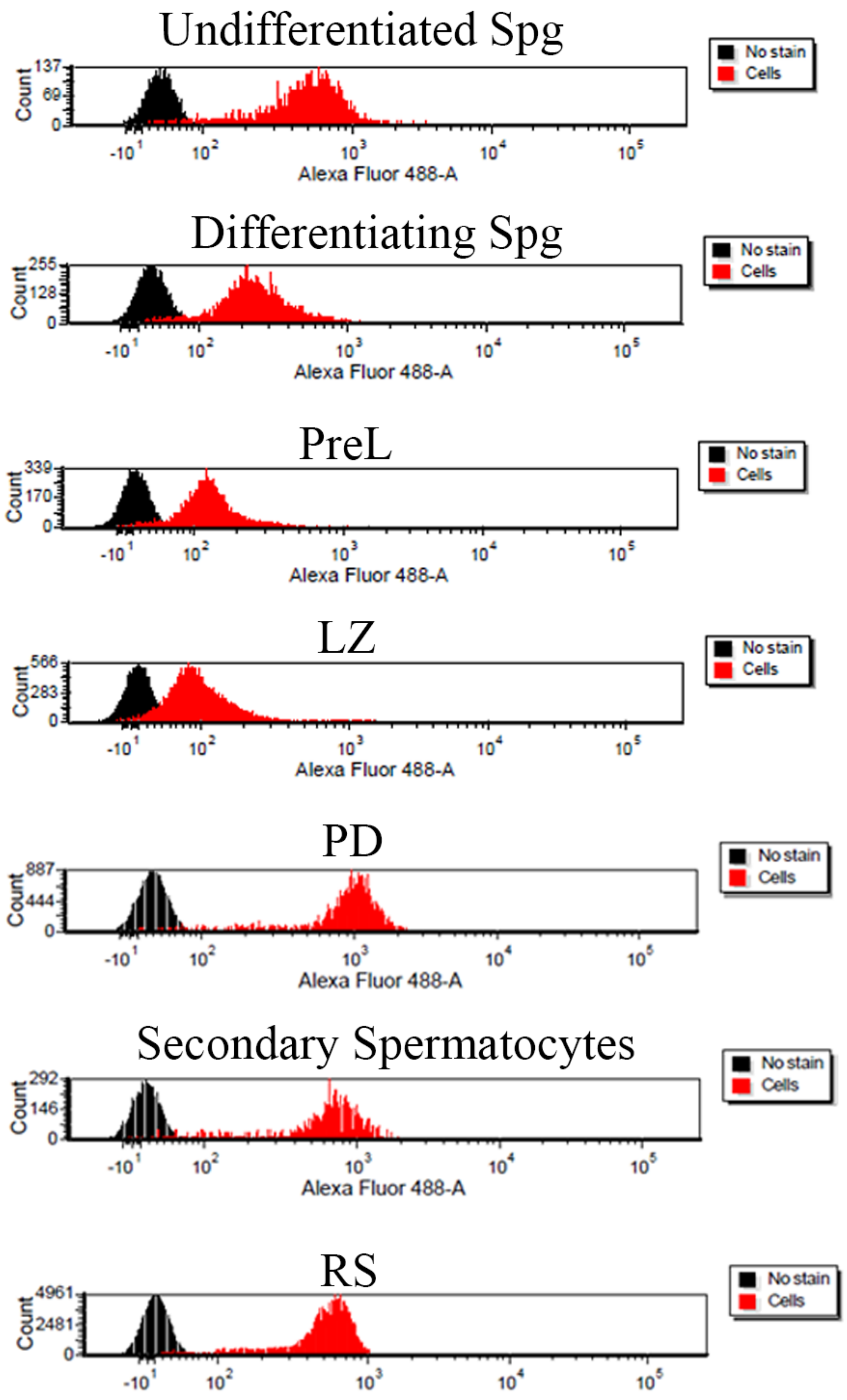
Undifferentiated Spg
Differentiating Spg
PreL
LZ
PD
Secondary Spermatocytes
RS
No stain
Cells
Count
Alexa Fluor 488-A

**Figure S2: Mitochondrial content dynamics across spermatogenesis**

Representative flow cytometry analysis (1 of 4 independent experiments) showing relative mitochondrial content per cell across spermatogenic populations, measured by MitoTracker Green staining. Shown populations include undifferentiated Spg, differentiating Spg, PreL, LZ, PD, secondary spermatocytes and RS. Histograms display fluorescence intensity for cell-containing samples (red) and background controls (black).

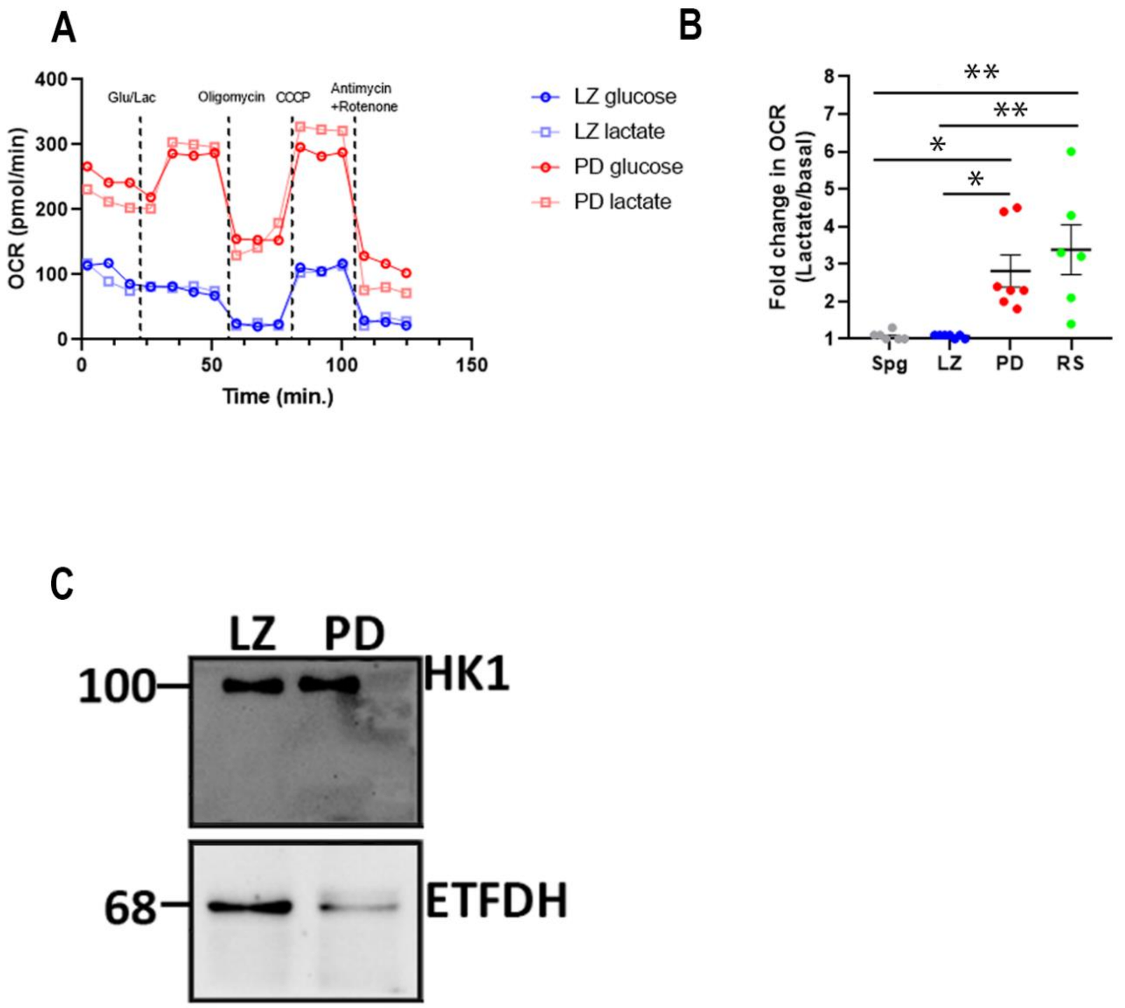


**Figure S3: Substrate-dependent respiration across spermatogenic stages**

(**A**) Representative OCR measurements of LZ and PD spermatocytes incubated in carbon-free basal buffer, followed by addition of 5 mM glucose or 5 mM lactate (Glu/Lac), and sequential injections of oligomycin 1µM, CCCP 5 µM and antimycin A/rotenone, each 1µM. (**B**) Quantification of lactate-induced OCR response (fold change relative to basal) in Spg, LZ, PD and RS; each point represents a biological replicate, bars indicate mean ± SEM and p-values are shown (ANOVA). *P < 0.05; **P < 0.01; ***P < 0.001; ****P < 0.0001. (**C**) Representative immunoblot analysis of ETFDH in LZ and PD cells, with HK1 as a loading control. Equal amounts of protein (40 µg) from each population were loaded onto separate gels.

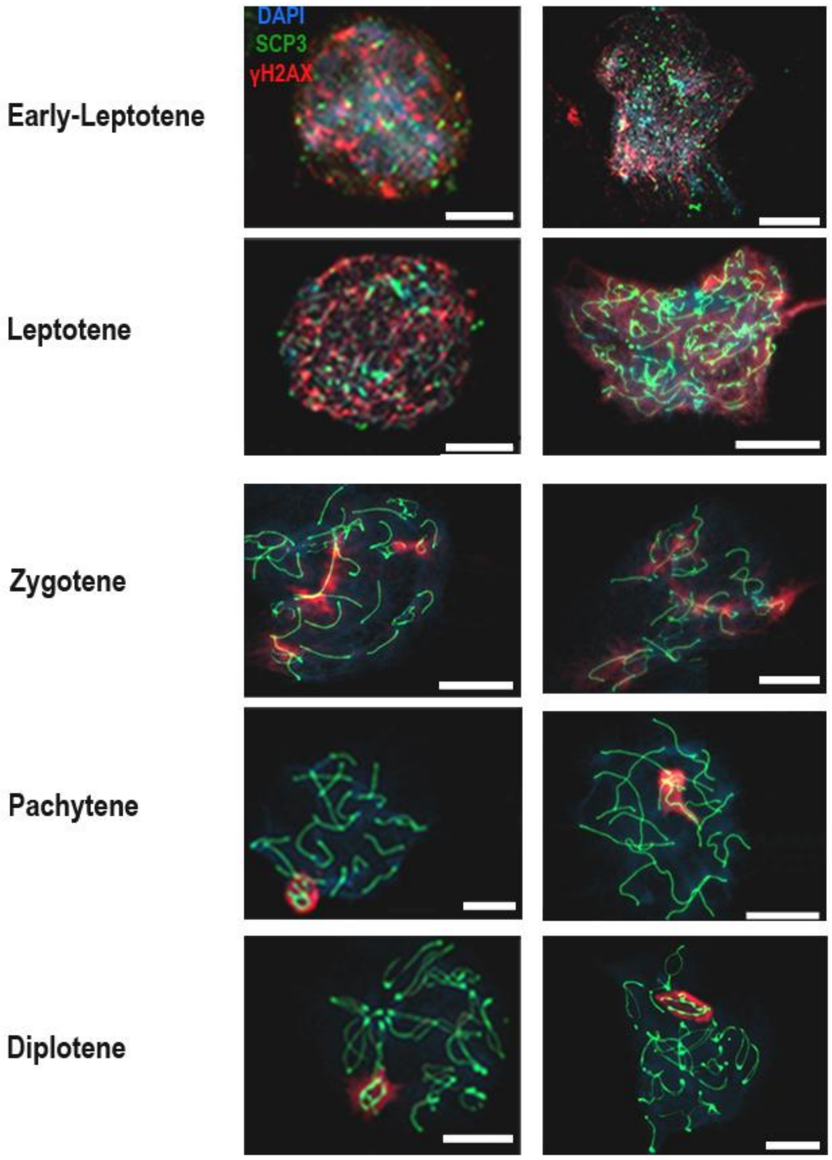


**Fig. S7. Nuclear spreads of spermatogenic cells from BRQ+uridine - treated mice**

Representative nuclear spreads of spermatogenic cells from BRQ+uridine - treated Stra8-tdTomato testes, stained for SCP3 (synaptonemal complex; green), γH2AX (DSBs/sex body; red), and DNA (DAPI; blue). Two representative cells are shown for each meiotic stage-early leptotene, leptotene, zygotene, pachytene and diplotene. Uridine supplementation restores normal nuclear morphology across all stages, with fully synapsed SCP3 tracks in pachytene and diplotene cells and γH2AX restricted to the sex body. Scale bars = 10 μm.

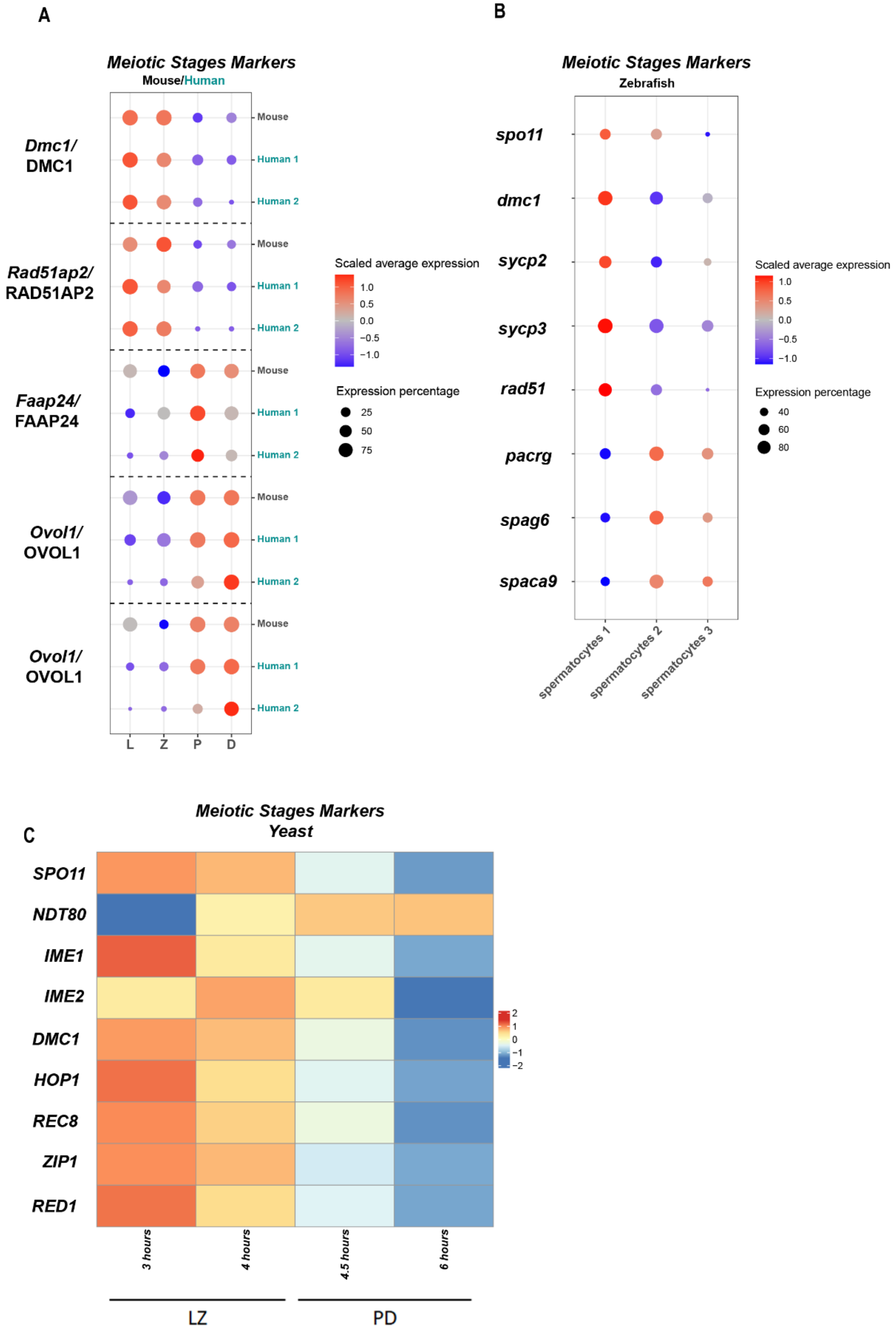

A
Meiotic Stages Markers
Mouse/Human
Dmc1/
DMC1
Rad51ap2/
RAD51AP2
Faap24/
FAAP24
Ovol1/
OVOL1
Ovol1/
OVOL1
Mouse
Human 1
Human 2
L
Z
P
D
Scaled average expression
1.0
0.5
0.0
−0.5
−1.0
Expression percentage
25
50
75
B
Meiotic Stages Markers
Zebrafish
spo11
dmc1
sycp2
sycp3
rad51
pacrg
spag6
spaca9
spermatocytes 1
spermatocytes 2
spermatocytes 3
Scaled average expression
1.0
0.5
0.0
−0.5
−1.0
Expression percentage
40
60
80
C
Meiotic Stages Markers
Yeast
SPO11
NDT80
IME1
IME2
DMC1
HOP1
REC8
ZIP1
RED1
3 hours
4 hours
4.5 hours
6 hours
LZ
PD
2
1
0
−1
−2

**Figure S9: Marker genes used to define early vs late meiotic prophase I populations across species.**

**(A)** Dot plots showing expression of prophase 1 substages marker genes in mouse and human single-cell RNA-seq datasets; dot color indicates scaled average expression and dot size indicates the fraction of expressing cells. **(B)** Dot plot of marker genes distinguishing zebrafish spermatocyte subpopulations (spermatocytes 1–3). Spermatocytes 1 roughly corresponds to LZ stages, spermatocytes 2-3 roughly correspond to PD stages. **(C)** Heat map of conserved meiotic-stage markers across a synchronous meiosis time course in S. cerevisiae, with early (LZ) and late (PD) time points indicated; colors represent scaled expression (z-score).
